# Novel Antibody Interfaces Revealed Through Structural Mining

**DOI:** 10.1101/2022.04.06.487381

**Authors:** Yizhou Yin, Matthew Romei, Kannan Sankar, Lipika R. Pal, Kam Hon Hoi, Yanli Yang, Brandon Leonard, Gladys De Leon Boenig, Nikit Kumar, Marissa Matsumoto, Jian Payandeh, Seth F. Harris, John Moult, Greg A. Lazar

## Abstract

Antibodies are fundamental effectors of humoral immunity, and have become a highly successful class of therapeutics. There is increasing evidence that antibodies utilize transient homotypic interactions to enhance function, and elucidation of such interactions can provide insights into their biology and new opportunities for their optimization as drugs. Yet the transitory nature of weak interactions makes them difficult to investigate. Capitalizing on their rich structural data and high conservation, we have characterized all the ways that antibody Fab regions interact crystallographically. This approach led to the discovery of previously unrealized interfaces between antibodies. While diverse interactions exist, β-sheet dimers and variable-constant elbow dimers are recurrent motifs. Disulfide engineering enabled interactions to be trapped and investigated structurally and functionally, providing experimental validation of the interfaces and illustrating their potential for optimization. This work provides first insight into previously undiscovered oligomeric interactions between antibodies, and enables new opportunities for their biotherapeutic optimization.

## Introduction

Avidity-driven amplification of weak transient protein-protein interactions is a common theme in immunological processes. In some instances, weak interactions are clustered at cell-to-cell synapses, e.g. between T cells and antigen presenting cells or target cells. In other cases, protein-level immune complexation can promote naturally weak monovalent affinities to stronger avidity-driven binding events, such as occurs during B cell receptor (BCR) selection and antibody responses (Lingwood et al., 2012).

While the antibody variable region (Fv) commonly binds target antigen with high affinity, monomeric interaction of the fragment crystallizable (Fc) region with effector receptors typically occurs in the µM range where 1:1 binding events are generally inconsequential. Yet within immune complexes, avidity amplifies these interactions into triggers for positive or negative cellular response. A recent illustration of biological selection for avidity-driven triggers is the discovery of a transient homomeric interface in the IgG Fc region that mediates antibody hexamerization (Diebolder et al., 2014). Nature’s ostensible purpose for this interface is the amplification of antibody-mediated complement pathways, which are initiated by interaction of the IgG Fc with pentameric complement protein C1q.

Apart from understanding the contribution of weak antibody interfaces to immunology, their further value is their utility for optimization. Monoclonal antibodies are the most successful class of biotherapeutics, delivering enormous impact for the treatment of cancer, autoimmunity, and other diseases (Carter and Lazar, 2018). As a consequence, antibodies have become one of the most highly engineered protein families. Beyond Fv affinity maturation for target binding, native IgG interfaces between immunoglobulin (Ig) domains and between antibody and cognate Fc receptors have been successfully optimized for enhanced activities. Examples include engineering of the IgG CH3 domain for heterodimerization (Ridgway et al., 1996) that enables bispecific antibody platforms (Spiess et al., 2015), enhancement of FcγR- and complement-mediated effector functions (Chu et al., 2008; Idusogie et al., 2001; Lazar et al., 2006; F. Mimoto et al., 2013; Futa Mimoto et al., 2013; Moore et al., 2010; Niwa et al., 2004; Richards et al., 2008; Shields et al., 2002, 2001; Stavenhagen et al., 2007; Umaña et al., 1999), and half-life extension through binding optimization to the neonatal Fc receptor FcRn (Dall’Acqua et al., 2006; Lee et al., 2019; Zalevsky et al., 2010). These types of subtle modifications to natural and sometimes weak antibody interfaces have met high success in drug development, with many of these platforms in clinically approved drugs (Gaudinski et al., 2018; Ghazi et al., 2011; Kolbeck et al., 2010; Latour et al., 2020; Nordstrom et al., 2011; Ollila et al., 2019; Robbie et al., 2013; Rugo et al., 2021; Salles et al., 2021; Sheridan et al., 2018; Tobinai et al., 2017; Vijayaraghavan et al., 2020). A corollary is that the discovery of new antibody interfaces creates new opportunities for biotherapeutic optimization.

An intriguing aspect of the Fc hexamer discovery is that initial insights were derived from crystal contacts within an IgG structure that had been deposited in the PDB many years prior (Diebolder et al., 2014; Saphire et al., 2001). Distinguishing true biological assemblies from artifactual crystal contacts is a long-studied problem in structural biology (Capitani et al., 2016). Additional functionally relevant interfaces have been identified from crystal packing, including a Fab homomeric interface (Tamada et al., 2015) and oligomer interfaces in plastocyanin (Crowley et al., 2008). These examples illustrate that crystal contacts can provide insights into natural biological interfaces, albeit in an ad-hoc manner.

Antibodies offer a unique opportunity to investigate interfaces in crystallographic data in an exhaustive manner. The high degree of structural homology of Ig domains together with the large number of antibody structures in the protein data bank (PDB) provide a rich dataset of interactions. In this study we have comprehensively mined the PDB to search for antibody interactions that may not have been previously realized. This work advances a novel structural informatic approach for the investigation of weak protein-protein interactions, and offers new insights into transient interactions in antibodies that may be relevant to immune biology and enable new capabilities for biotherapeutic optimization.

## Results

### Computational pipeline identifies common interactions from Fab-Fab crystal packing

To search for new antibody interfaces, we computationally analyzed the crystal contacts for all Fab structures in the PDB (***Figure 1A***). We leveraged the structural antibody database SAbDab (Dunbar et al., 2014), which at the time of our experiments totaled 1,456 Fabs from X-ray diffraction. The structures included apo Fabs as well as those in complex with target antigen, and both were included in the calculations.

**Figure 1.**
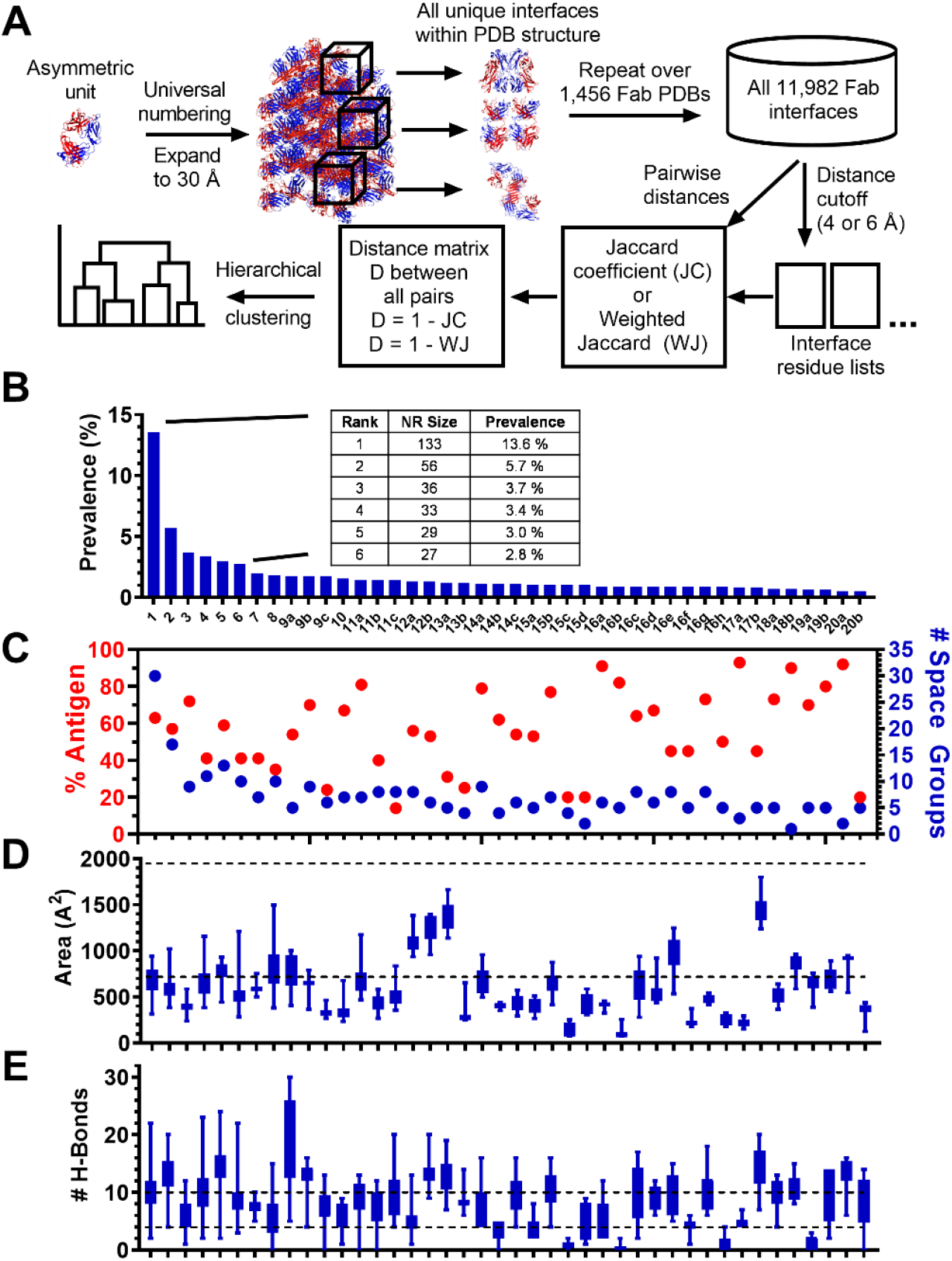
Antibody Fab contacts and Fab-Fab interfaces revealed from informatic PDB analysis. (**A**) Computational flow diagram for analyzing inter-Fab interfaces. Fab X-ray structures were universally renumbered and the asymmetric unit was expanded to 30 Å^2^. Symmetric oligomers were extracted, and all interacting residue pairs were identified using a distance cutoff of either 4 or 6 Å between any two inter-Fab residues. Similarity between a pair of interfaces was calculated as the Jaccard index or weighted Jaccard index between the sets of interface residue positions within each interface. Hierarchical clustering was performed based on the similarities calculated to result in a dendrogram of Fab interfaces. (**B**) Recurrent packing Fab-Fab interfaces throughout the collective Fab PDB in order of decreasing prevalence. Prevalence reflects an incidence measure of each interface that is unbiased by the presence of multiple structures of the same Fab sequence in the PDB. The inset provides the nonredundant (NR) size and prevalence values for the 6-most prevalent clusters. (**C**) Percentage of PDBs within each of the 42 most prevalent clusters that include antigen in the structure (red, left axis), and number of distinct space groups among the PDBs within each cluster (blue, right axis). The plot shows that the lack of cluster bias due the presence of antigen or crystallographic lattice. For example, the six most prevalent interfaces were observed in structures where Fab/antigen complexes make up 63%, 57%, 72%, 41%, 59%, and 41% of the cluster respectively, and that crystallized in 30, 17, 9, 11, 13, and 10 different space groups respectively. (**D**) Mean buried surface area and (**E**) number of H-bonds for the 42 most prevalent interfaces, in order of decreasing prevalence. Error bars represent standard deviations. Data in **B-E** are vertically aligned and thus there is correspondence with labeled columns in (**B**). Numeric values for prevalence and PISA results are provided in ***Supplementary table 1***.

Structural analysis of Fabs is facilitated by their high degree of structural and sequence homology; a position in any given Fab is generally structurally equivalent to that same position in virtually all other Fab structures. To enable comparisons, variable heavy (VH) and light (VL) regions were renumbered according to the AHo numbering convention (Honegger and Plückthun, 2001), which accounts for sequence length differences to accurately represent structural equivalence. Because the commonly used EU convention (Kabat et al., 1991) for constant regions does not account for gaps across diverse Ig sequence space, we developed a custom constant region numbering convention (Materials and Methods). While our computational methods used these more robust conventions, for literature consistency this manuscript labels variable and constant region positions according to Kabat and EU numbering, respectively.

A computational pipeline was built to analyze the crystal contacts for all Fab structures (***Figure 1A***). Fab asymmetric units were expanded, all non-redundant symmetric oligomers within the crystal lattice were identified, and non-redundant interfaces in these oligomers were extracted. 11,982 total interfaces were identified across 1,456 PDB entries analyzed. All inter-Fab residue pairs constituting the interfaces were identified using a 4 or 6 Å distance cutoff. An inter-Fab contact map (***Supplementary figure 1***) revealed patterns both on and off the diagonal, representing homodimeric and heterodimeric interactions respectively.

### Antibody Fabs oligomerize through common interfaces

Cluster analysis was performed on the 11,982 interfaces to assess recurrence (***Figure 1A***). Similarity between all interfaces was calculated in a pairwise manner as the Jaccard index between the sets of interface positions within each interface, followed by hierarchical clustering of similarity values (Materials and Methods). From the analysis it was evident that the majority of interface clusters were singletons, i.e. they occur only once in the analyzed set of PDBs. The focus of this study was on interfaces that occur frequently across multiple PDB structures.

The results revealed recurrent Fab packing interfaces. Total PDB cluster size (***Supplementary table 1***) represents the number of distinct Fab PDBs from the 1,456 set that contain a given interface at least once in the lattice expansion. However, the PDB set contains structures with identical Fab sequences, and thus the total size for each cluster includes redundant Fabs. Identical sequences (%ID=100) within each cluster were reduced, leaving 981 nonredundant (NR) Fabs from the original set of 1,456. NR size divided by 981 nonredundant sequences is referred to as prevalence (***Figure 1A***). The most recurrent interface cluster occurs 133 times in the 981 nonredundant Fab PDB structures for a prevalence of roughly 14%. The next 5 most recurring interfaces occur with a prevalence of 5.7%, 3.7%, 3.4%, 3.0%, and 2.8% respectively (***Figure 1B and Supplementary table 1***). The Fab PDB comprises apo-Fab structures as well as Fab-antigen complexes. Fab-antigen contacts were not part of the calculations, and there was no apparent cluster bias due to the presence or absence of antigen (***Figure 1C***). These results suggest not only that the observed interfaces are uninfluenced by antigen, but further that they are robust across structures of Fab complexes with diverse target antigens. Finally, cluster members crystalized in a diversity of space groups (***Figure 1C***), suggesting no bias of interfaces by crystallographic lattice.

The observed Fab dimers generally had structural and energetic features of weak transient interfaces, as analyzed using the ‘Protein interfaces, surfaces and assemblies’ service PISA (Krissinel and Henrick, 2007). With some exceptions, the generally lower buried surfaces of the discovered interfaces (562 ± 314 Å^2^) (***Figure 1D***) are more similar to “weak transient complexes” (718 ± 195 Å^2^) than “obligate” or “permanent homodimers” (1,950 ± 986 Å^2^) (Luo et al., 2015). In contrast, the numbers of Fab interface H-bonds (8 ± 4) (***Figure 1E***) are closer to obligate homodimers (10 ± 8) than transient complexes (4 ± 3) (Luo et al., 2015). The more robust H-bonding relative to buried surface may be skewed in part by the prevalence of β-sheet dimers, discussed further below.

Each cluster is a distinct Fab-Fab interface. We established a naming convention based on the structurally central residue of each discovered interface (Materials and Methods). For this convention, the name of each cluster adopts the format X-#, in which X represents the chain (VH, CH1, VL, CL) and # represents the position of the residue at the structural center of the interface, numbered by Kabat for VH and VL or EU for CH1 and CL. In addition, we designated a structural representative for each cluster, defined as the member with the highest weighted Jaccard similarity index to all other members in the cluster (Materials and Methods). ***Figure 2*** presents the representative Fab dimer for each cluster with a prevalence of ≥ 0.5%. The interfaces, 42 total, are ranked numerically based on prevalence, with equivalent prevalence designated with arbitrary alphabetic qualifiers. The labels within ***Figure 2*** include the cluster rank, interface name, prevalence, and buried surface area (described below). Interface positions for all of the top 42 clusters are provided in ***Supplementary table 2***. Pymol files for all 42 interfaces will be provided in the supplementary materials.

**Figure 2.**
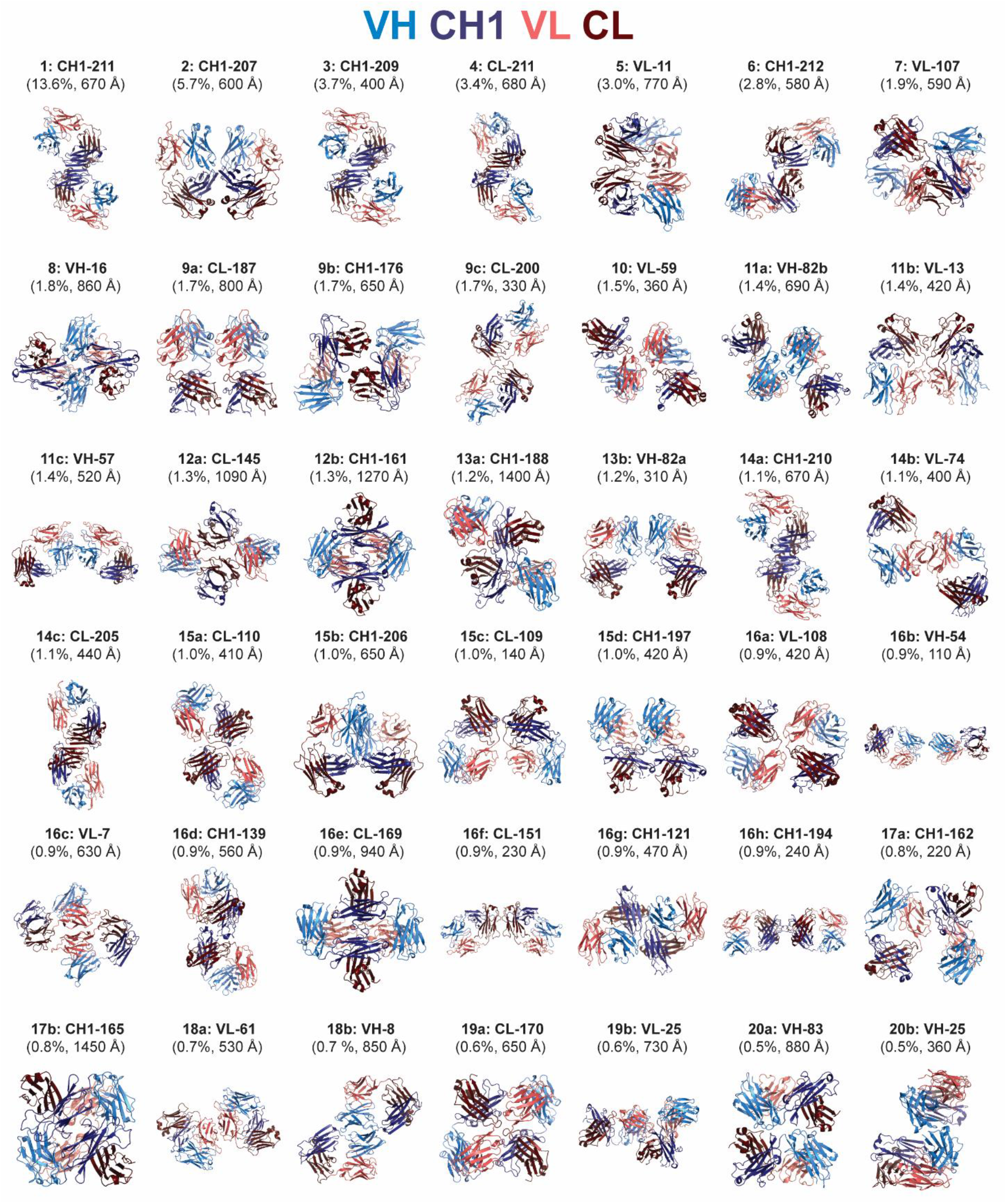
Representative structures of the 42 most prevalent Fab-Fab interfaces. Domains are colored as follows: VH (light blue), CH1 (dark blue), VL (pink), CL (dark red). The labels for each interface include the cluster rank based on prevalence, interface name based on structurally central residue (Kabat numbering for Fv region, EU number for constant regions), nonredundant prevalence percentage, and mean buried surface area. Interfaces are ranked numerically based on prevalence, with equivalent prevalence being designated with arbitrary alphabetic qualifiers; for example, there are three interfaces that occur in 17 PDB entries for prevalence = 1.7%, ranked as 9a, 9b, and 9c.

### β-sheet dimers and elbow dimers are common Fab interface motifs

The most prevalent interface, labeled CH1-211, is a sheet-extended homodimer mediated through the heavy chain (HC) constant region centered on (EU) position 211 (***Figure 2***). This interface is observed in 133 of nonredundant Fab PDB structures, or 13.6% of the nonredundant Fabs (***Figure 1B***). The extended strand pairing is mediated centrally by the C-terminal strand G (positions 207-214) that effectively doubles the size of the β-sheet. In addition, the interface is complemented by packing of the short helical turn of CL between strands A and B (122-126) against the N-terminus of CH1 prior to strand A (119-122). Altogether this interface buries an average 670 Å of surface calculated across the members of the cluster.

Strikingly, β-sheet dimers were found to be a common interaction motif, observed in a total of eight clusters (***Figure 3A***). Two additional CH1 homodimer interactions are observed as interfaces CH1-209 and CH1-210, which correspond to the 3rd and 14th (14a) most prevalent clusters (3.7% and 1.1% respectively). These dimers are highly similar to CH1-211, containing the same sheet-extended HC homodimer yet with either a 2-residue (CH1-209) or 1-residue (CH1-210) register shift. While interface CH1-210 contains similar LC packing arrangement, the greater register shift of CH1-209 separates the dimer units such as to eliminate the LC contacts, resulting in less buried surface (400 Å averaged over the cluster) (***Figure 3A***). In addition to CH1 homodimers, sheet-extension interfaces make up two CH1/CL heterodimers (CL-211 and CH1-212), a CL homodimer (CL-205), a VH homodimer (VH-57), and a VL homodimer (VL-11). Altogether, β-sheet dimers make up five of the six most prevalent interfaces, and appear in over 30% of nonredundant PDB entries. The structural similarity of the members within interface clusters CH1-211, CL-211, and CL-205 are high (***Figure 3B***), further illustrating the common structural motif. A high degree of structural homology was observed upon superposition of the structural representatives of the CH1-211 homodimer with the CL-211 heterodimer and CL-205 homodimer (***Figure 3C***).

**Figure 3.**
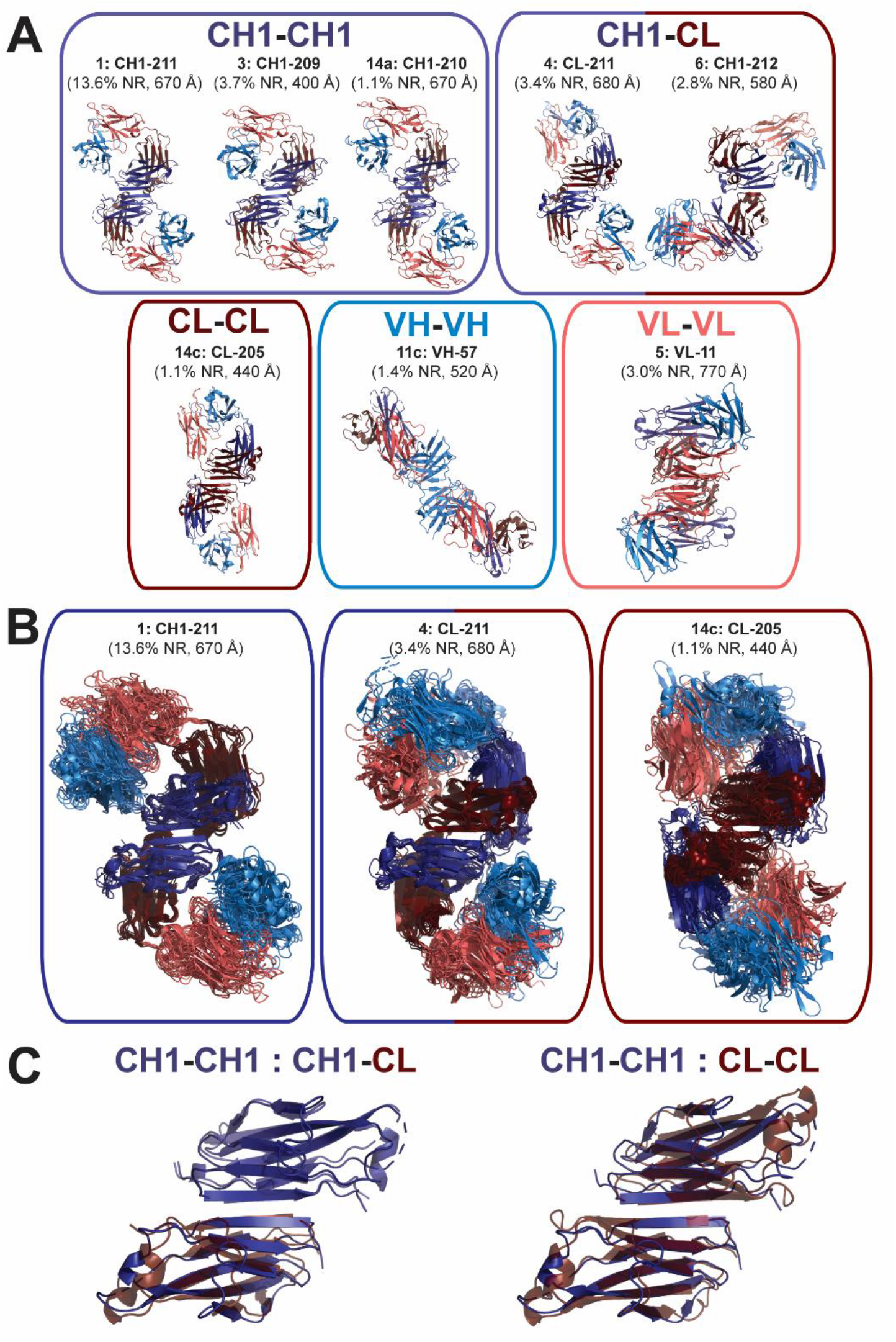
β-sheet dimers are commonly observed Fab oligomers. (**A**) Structural similarity between sheet-extended Fab oligomers that interact as CH1-CH1 homodimers (top left), CH1-CL heterodimers (top right), CL-CL homodimers (bottom left), VH-VH homodimers (bottom middle), and VL-VL homodimers (bottom right). Domains are colored as follows: VH (light blue), CH1 (dark blue), VL (pink), CL (dark red). The labels for each interface include the cluster rank based on prevalence, interface name based on structurally central residue, nonredundant prevalence percentage, and mean buried surface area. (**B**) Fab β-sheet dimers are structurally similar across cluster members. Interface residues of 10 cluster members were superimposed for Fab oligomers within the CH1-211 homodimer (left), CL-211 heterodimer (middle), and CL-205 homodimer (right). (**C**) Constant domain superposition of CH1-CH1 homodimer with CH1-CL heterodimer (left) and CL-CL homodimer (right). β-sheet residues at the Fab-Fab dimer interface were superimposed for the structural representatives of clusters CH1-211 and CL-211 (left), or CH1-211 and CL-205 (right).

All of the β-sheet dimers position the Fab monomers in an antiparallel orientation, with opposed Fv’s. In the context of a full-length antibody, using the most N-terminal hinge disulfide (EU position Cys226 in human IgG1) as the tether point and a random coil residue distance of 3.5 Å, the C-termini of the two Fab arms are constrained to a maximal distance of ∼35 Å relative to each other. The calculated distance between the HC C-termini of the two Fabs in all of the β-sheet dimers exceed this value (CH1-211: 41 Å, CH1-209: 47 Å, CH1-210: 37 Å, CL-211: 47 Å, CH1-212: 37 Å, CL-205: 43 Å). This analysis suggests that the orientation of the two Fabs precludes intra-IgG binding, forcing inter-IgG interactions that would ostensibly promote higher-order complexation.

Beyond β-sheet dimers, an additional recurring motif observed are interfaces mediated by the elbow regions between variable and constant domains (***Figure 2 and Supplementary figure 2***). These interactions involve the elbow regions between either two HCs (CH1-207 and CH1-121) or two LCs (CL-200, CL-110, VL-108, CL-109, CL-169, and CL-170). CH1-207 is the second most commonly observed interface, and in total, the eight elbow region interfaces comprise 12.7% of nonredundant PDB entries. The proximity of both the elbow and the β-turns/loops varies among these eight interfaces. In the case of CH1-207, there are roughly balanced contributions from the variable and constant region loops and turns (***Supplementary table 2***). While elbow-elbow interaction was the general theme among these clusters, as a collection there was greater overall structural dissimilarity relative to the group of β-sheet dimers (***Supplementary figure 2***).

### Intra-cluster members show similarities at interface residues

To investigate sequence dependence, interface profiles were generated for the six most prevalent clusters (***Figure 4***), as well as the three additional clusters that form β-sheet dimers (VH-57, CH1-210, and CL-205) (***Supplementary figure 3***). The profiles describe the compositional distribution of each cluster based on species (human, mouse, other), light chain (LC) type (kappa or lambda), as well as human and mouse VH and VL subgroups. In addition, the profiles provide intra-versus inter-cluster sequence identity for both the entire Fv as well as only those residues at each corresponding interface. For the Fv, these values reflect the mean pairwise identities for all VH and VL sequences within a cluster (intra) versus across the clusters (inter). A similar comparison is made for the interface, where only those residues that participate in the interface for the designated cluster are compared. Finally, sequence logos provide weighted sequence composition for cluster members at interface residues.

**Figure 4.**
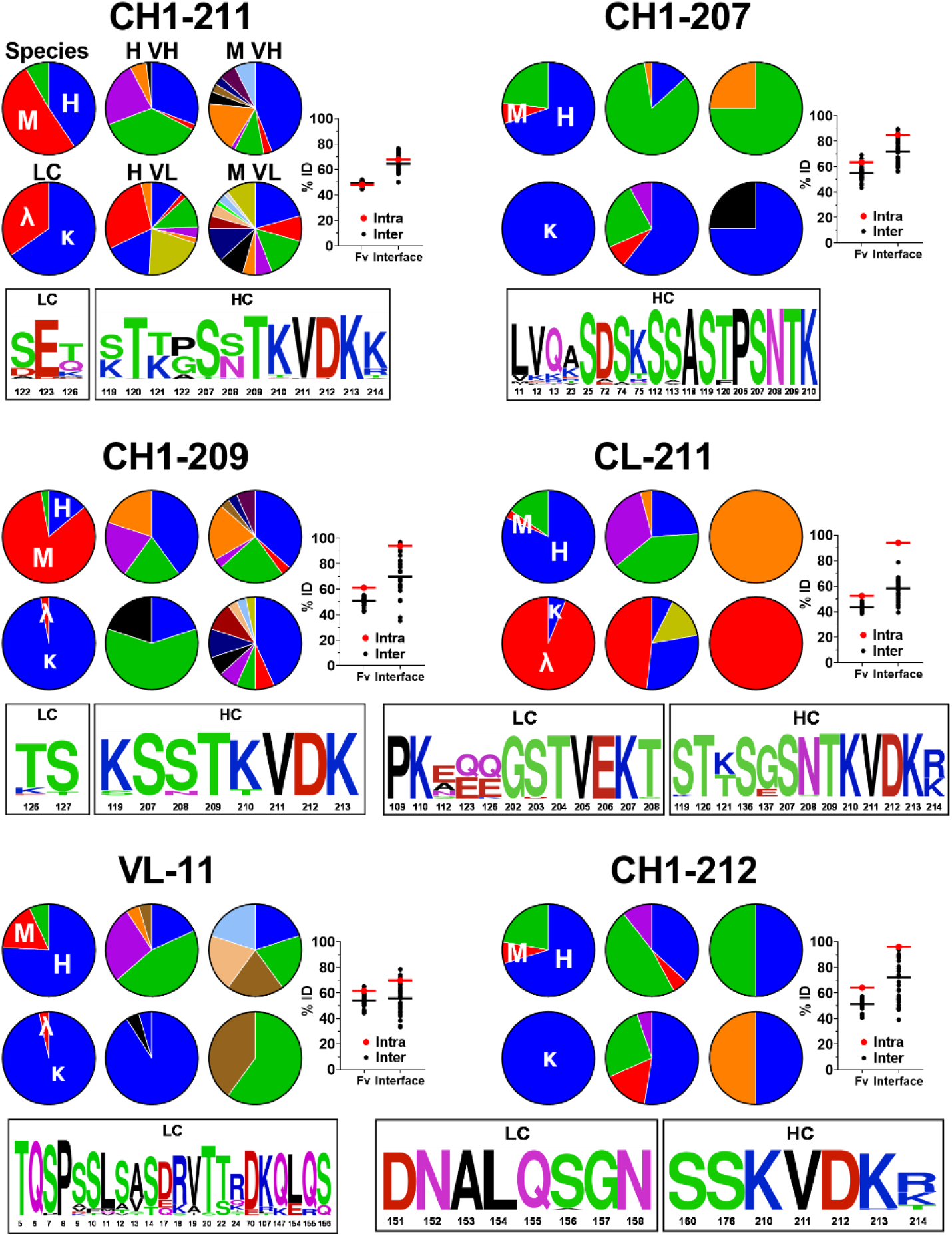
Interface profiles of the six most prevalent clusters. For all interfaces, the upper left-most pie chart provides the sequence distribution of cluster members based on species: human (blue, H), mouse (red, M), and other (e.g., rat, rabbit, etc., green). The lower left-most pie chart provides the sequence distribution of cluster members based on light chain type: kappa (blue, κ), and lambda (red, λ). The four right-most pie charts provide the sequence distribution of human VH (H VH), human VL (H VL), mouse VH (M VH), and mouse VL (M VL) subgroups. The labels above the pie charts in the upper left CH1-211 profile are not repeated in the other charts for visual simplicity. The %ID plot on the right provides the intra (red) versus inter (black) cluster sequence identity for both the entire variable region (Fv) as well as only those residues at the interface as shown in the sequence logo and ***Supplementary table 2***. For the Fv, these values reflect the mean pairwise identities for all VH and VL sequences within the cluster (intra) versus the mean pairwise identities between each member of the cluster and all other members of all other clusters (inter). A similar comparison is made for each interface, where %ID reflects the mean pairwise identity of the residues at the interface of a given cluster aligned with those same residues for each member of the cluster (intra) versus all other members of all other clusters (inter). The sequence logo at the bottom provides weighted sequence composition at interface residues for members of the designated cluster, with numbering according to Kabat and EU conventions for the Fv and constant regions, respectively.

CH1-211 showed no apparent species dependence, being represented by sequences from diverse species, LC type, and VH and VL subgroups (***Figure 4***). The diverse subgroup representation is also captured in the similar Fv intra-vs. inter-cluster identities (∼50%). Similar overall results were observed for the other top six clusters, with most demonstrating diverse representation of species, LC, and Fv subgroups (***Figure 4***). Notable biases are the Fv subgroup trends of CH1-207 and to lesser extent VL-11. These are the two clusters among the top six where the Fv contributes substantially to the interface, specifically the VH for the former and VL for the latter (***Figure 2***). These clusters are comprised principally of human subgroup IGHV3 for CH1-207 and human subgroup IGKV1 for VL-11. The consensus interface residues based on their sequence logos (***Figure 4***) are well-represented in these germlines.

The interface residues of CH1-211 are roughly as similar among members of the cluster (intra identity 68%) as they are to those same residues among members of all other clusters (inter identity 64%) (***Figure 4***). In contrast, three of the top clusters, CH1-209, CL-211, and CH1-212, display high intra-cluster interface identity (94%, 94%, and 96% respectively) relative to lower inter-cluster identities (70%, 58%, and 72% respectively) (***Figure 4***). CL-211 is notable also for its bias towards lambda LCs (***Figure 4***). While kappa LCs are overrepresented in the PDB set overall (roughly 85% are kappa, 15% are lambda), nearly all of the members of CL-211 are lambda. Deeper investigation revealed that kappa LCs contain a proline at position 204 that terminates the N-terminus of the last β-strand. In contrast, lambda LCs contain a Thr at 204 that, together with a shorter N-terminal loop, enable an extended β-strand that permits β-sheet heterodimerization with the HC. Consistent with this observation, the CL-CL homodimer interface CL-205 also showed a bias for lambda (***Supplementary figure 3***). Overall, the diverse nature of the sequences at a global level yet trends for some clusters at an interface level, suggest determinant motifs that may provide further insight into the interactions as well as serve as guides for design.

### Disulfide engineering enabled Fab dimers to be trapped

While some of the observed interfaces may be artifacts of crystallization, we experimentally investigated the prospect that some may reflect natural albeit transient Fab oligomeric interactions in solution. Given their weak nature, we explored whether dimers could be trapped via engineered disulfides. Starting from the representative structures of each of the 42 most prevalent clusters (***Figure 2***) we performed in silico cysteine scanning analysis to explore all possible residue pairs at each interface for distance and geometry that may be conducive to disulfide bond formation (Materials and Methods). A total of 179 residue pairs were identified, spanning 77 HC-HC, 54 LC-LC, and 48 HC-LC pairs. While the range of designed variants per cluster was as high as 14, the median was 2 variants per cluster. Cysteine variants were engineered into the Fab region of the anti-Her2 antibody trastuzumab (Carter et al., 1992). Variant and WT Fab antibodies were expressed in HEK 293 cells, purified by affinity resin, and analyzed by size exclusion chromatography (SEC) (***Figure 5A***). Analysis of the full set of analytical data revealed that many variants showed discrete dimeric species, in addition to monomeric antibody (***Figure 5B and Supplementary tables 3-5***).

**Figure 5.**
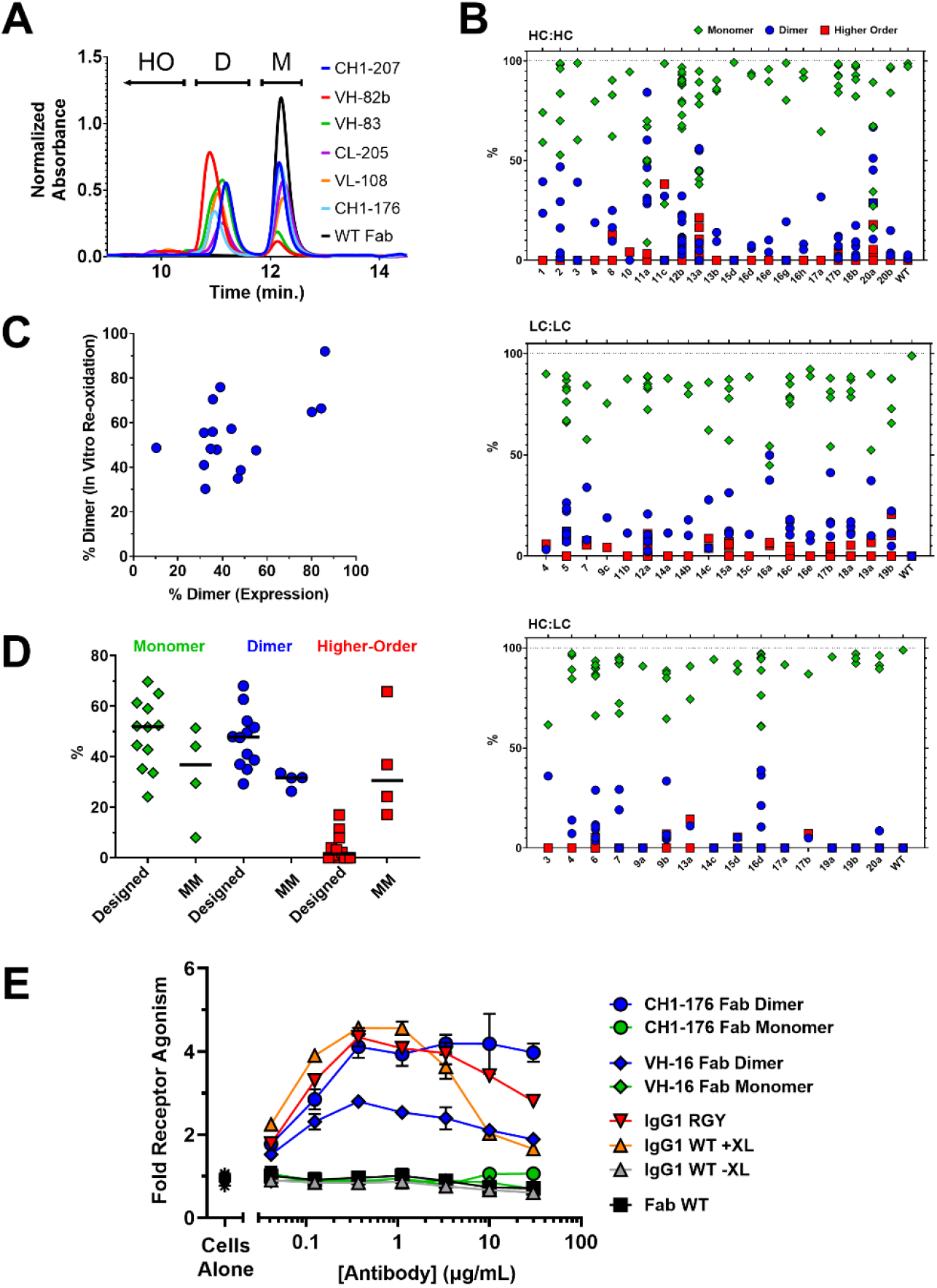
Fab dimers can be trapped with engineered disulfides. (**A**) Representative SEC chromatograms of select cysteine variant and WT anti-Her2 Fabs after affinity chromatography. Peaks are labeled as monomer (M), dimer (D), and higher-order (HO). (**B**) Summary of monomer (green diamonds), dimer (blue circles), and higher order (red squares) species for all cysteine variants by SEC post-affinity chromatography. Data represent percentage of species based on integration of SEC peaks. Multiple symbols for a given cluster represent multiple designed cysteine variants. The identities of all variants and associated numeric SEC data are provided in ***Supplementary tables 3-5***. (**C**) Correlation between expression and in vitro coupling data for disulfide-trapped dimers. (**D**) Summary of data from in vitro coupling of selected cysteine variant Fabs. The data reflect the % monomer (green diamonds), dimer (blue circles), and higher-order (red squares) species after assembly for the designed and mismatched (MM) variants run as negative controls. For MM variants the pairings of HCs and LCs with single cysteine mutations were scrambled in four separate combinations. The identities of all variants and associated numeric data are provided in ***Supplementary table 6***. (**E**) OX40 receptor activation by variant and WT Fab and full-length IgG antibodies based on an NFκB luciferase reporter assay. VH-16 represents Fab cysteine variant VH(S113C) / CH1(G178C) and CH1-176 represents Fab cysteine variant VH(P14C) / CL(D151C). Fold receptor activation represents normalized RLU relative to cells alone. For the IgG1 versions, +XL and -XL indicate the presence or absence of secondary crosslinker antibody, respectively. RGY represents a triple Fc variant that promotes IgG hexamer formation. No crosslinker was used for the RGY version or the Fab monomer or dimer versions.

The presence of discrete dimer species in many of the designed variants supports the computational analysis. Yet many of the cysteine variants were >95% monomeric, which is a common expression profile for Fabs without engineered disulfides, as evidenced by WT (***Figure 5A and B***). These monomeric constructs represent internal controls, suggesting that transient non-random contact in solution as well as proper geometry are required to form the stable oligomeric species. A subset of variants with the most prominent dimer formation were selected to explore whether dimers could be assembled in vitro. The monomeric Fab species of this variant subset were reduced in solution followed by gradual re-oxidation with dehydroascorbic acid. Upon re-oxidation, discrete dimer species were formed, similar to and correlated with the results from in vivo expression (***Figure 5C and Supplementary table 6***). As a control, the pairings of HCs and LCs with single cysteine mutations were scrambled in four separate combinations, referred to a mismatched (MM) variants (***Supplementary table 6***). Upon refolding, all mismatched combinations displayed substantial higher order oligomers that indicate nonspecific disulfide formation (***Figure 5D and Supplementary table 6***). These results suggest that for the designs, ordered contact influences Fabs toward stable dimer species in solution, further supporting the presence of specific albeit transient interactions.

### Fab dimers potentiate antibody functional activity

A representative set of 26 cys-engineered variants were selected to explore functional application. Cysteines for each variant were introduced into an antibody referred to as 3C8 that is an agonist of the receptor OX40 (CD134), which we have previously shown to provide a sensitive system for detecting antibody-mediated receptor clustering (Yang et al., 2019). Due to low expression yields or variable monomer/dimer ratios, many of the variants were not characterized further. A subset of 5 variants that expressed well and had a favorable monomer/dimer profile were chosen for further purification and separation of species. Final monomer and dimer samples for each variant had high purity and were stable in solution (i.e, they did not interconvert over time based on analytical SEC). Affinity measurements by Biacore indicated that a subset of dimeric versions did not retain binding (data not shown), possibly due to steric clash of antigen binding in the context of the coupled Fab dimer. However, two variants, VH(S113C)/CH1(G178C) at the VH-16 interface and VH(P14C)/CL(D151C) at the CH1-176 interface, maintained their affinities for OX40 (K_D_’s for variants and WT ∼10 nM, ***Supplementary figure 4***).

In an NFκB luciferase reporter assay utilizing OX40+ Jurkat cells, the 3C8 antibody has no activity as a bivalent IgG on its own, but is a strong agonist of OX40 signaling when extrinsically crosslinked with a secondary antibody (***Figure 5E***). An Fc-engineered triple variant (RGY) of this IgG that promotes hexamerization (Diebolder et al., 2014) agonizes receptor without reliance on extrinsic crosslinker. Strikingly, Fab dimer versions of the two cysteine-linked variants were capable of agonizing OX40 in the absence of crosslinking, in contrast to inactive monomer versions of the same variants (***Figure 5E***). The ability of the cys-linked Fab dimers to activate receptor signaling despite their bivalency may be due to their ability to engage receptor with a particular geometry and/or orientation. While the mechanism requires further study, these results illustrate how new interfaces may be used to engineer novel geometries and orientations into antibodies in order to enable activities of therapeutic relevance.

### Disulfide-trapped Fab dimers are structurally similar to informatically-mined conformations

To validate proper disulfide trapping of the interfaces and further support the results, we solved the x-ray structures at <3 Å resolution for a representative subset of disulfide-trapped Fab dimers containing the trastuzumab Fv. Experimental structures included one representative of the β-sheet dimer class [CL-205 (***Figure 6A***), and two representatives of the elbow dimer class CH-207 (***Figure 6B***) and VL-108 (***Figure 6C***). Electron density maps were consistent with the presence of engineered disulfide bonds at the expected sequence positions (***Supplementary figure 5***). Experimentally-determined and informatically-mined structures were topologically similar, with good superposition at the interface sites (***Figure 6***). The most notable alignment discrepancy across the three can be seen for CH1-207. In this structure, the disulfide bond pulls the HC elbow loops closer together on one end. While this perturbation rotates the left Fab with respect to the right Fab, the interface region remains largely intact, suggesting that this discrepancy may be due to disulfide bond constraints rather than the formation of an altered solution-phase configuration. Overall, these results provide strong evidence that the observed solution-phase dimers homogeneously resemble the discovered dimer conformation rather than a mixture of interface-independent conformations from non-specific disulfide pairing.

**Figure 6.**
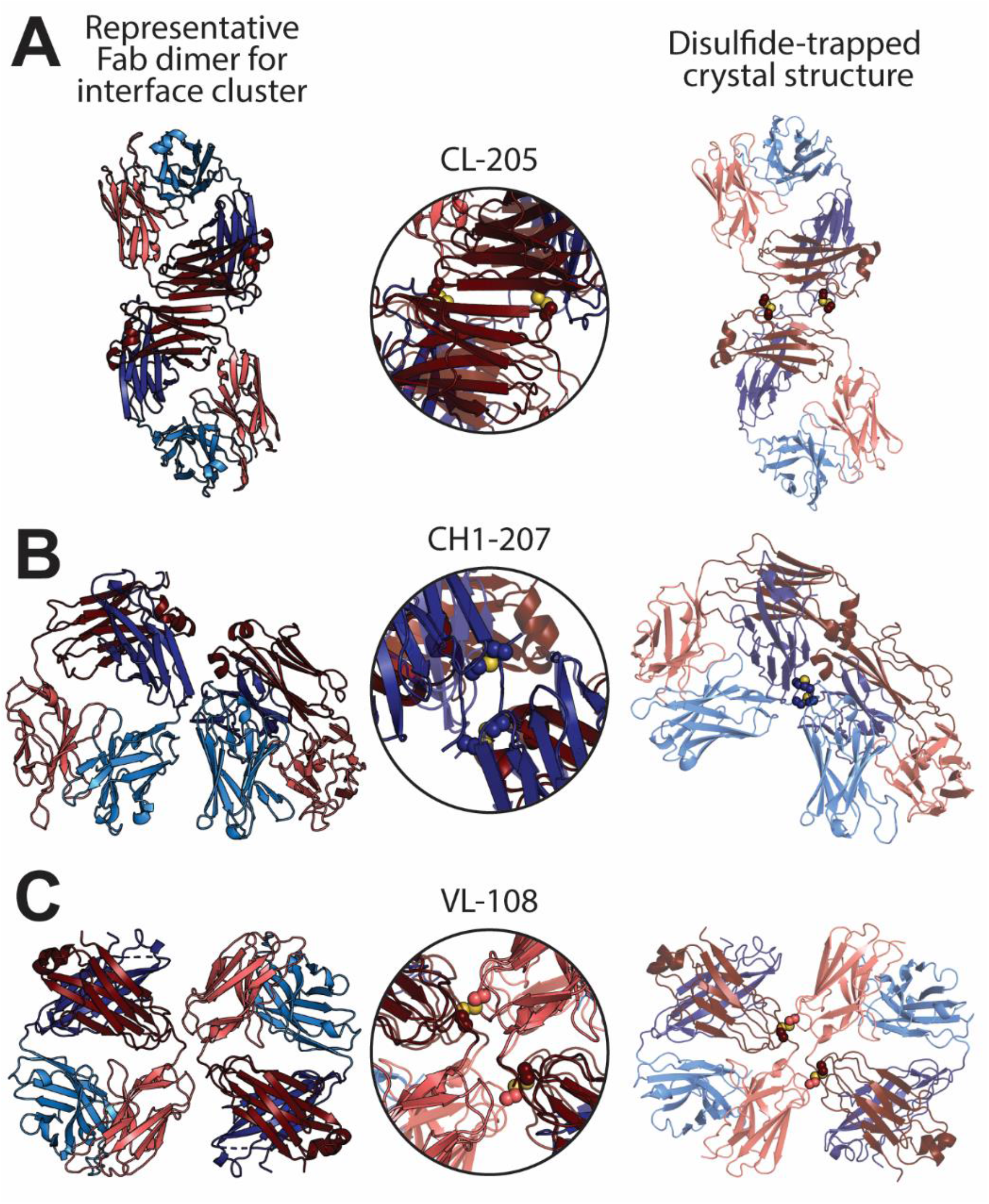
Disulfide-trapped Fab dimers affirm conformations of PDB mining. All Fabs comprise the variable region of the anti-Her2 antibody trastuzumab. Fab structures represent the β-sheet-extended interface cluster CL-205 [variant CL(S202C) / CL(S208C)] (**A**) and the elbow region clusters CH1-207 [variant CH1(S119C) / CH1(G122C)] (**B**) and VL-108 [variant VL(G16C) / CL(D170C)] (**C**). Fab domain colors match those in ***Figure 2***, with the informatically-mined structure at 0% transparency (left) and the experimentally-determined structure at 30% transparency (right). Sulfur atoms participating in disulfide bonds are shown in yellow. Overlays of the interface (middle) for **A** and **C** were configured by aligning the α carbons of the Fab dimers from each structure, whereas the overlay for **B** used a single Fab for alignment (see Results).

## Discussion

Weak transient protein-protein interactions play fundamental roles in diverse biological processes (Garcia-Seisdedos et al., 2017; Humphris and Kortemme, 2007; Ozbabacan et al., 2011). While best characterized in the context of intracellular signaling cascades, transient interactions are also relevant extracellularly, including during immunological recognition where they can convert the avidity of a cell synapse or antibody complex into a cooperative trigger for cellular activation or inhibition. While obligate interactions are readily investigated with direct biochemical and structural methods such as crystallography and electron microscopy, transient interactions are low in abundance and harder to identify, requiring sensitive and sometimes indirect techniques such as the yeast two-hybrid system, fluorescence resonance energy transfer, nuclear magnetic resonance spectroscopy, and split protein complementation (Ozbabacan et al., 2011; Romei and Boxer, 2013). While in rare instances weak interactions have been gleaned from crystallographic structures, it can be difficult to distinguish between true biological interfaces and crystal packing artifacts (Capitani et al., 2016).

We have explored a novel structural informatics approach to search for previously uncharacterized interfaces in antibodies. We selected antibodies for three reasons. First, the avid nature of immune complexation provides a biological rationale for the existence of as yet undescribed interfaces that could tune immune response. Second, the virtual one-to-one residue equivalence of antibodies across a large structural data set enables mining of interaction patterns for commonality. Effectively, prevalence in the current work serves as a signal-to-noise parameter that suggests biological relevance. Finally, we are interested in discovering new antibody interfaces for their potential in biotherapeutic engineering. Monoclonal antibodies are one of the most successful classes of drugs across a myriad of medical needs, with the 100th antibody drug recently approved (Mullard, 2021). While the most effective antibody drugs have historically been native IgGs, recent years have witnessed an acceleration in development and approval of engineered versions optimized for activity (Carter and Lazar, 2018). Rather than de novo sites, the most clinically successful enhancements are modest mutational modifications of natural and often weak interactions, either intra-IgG or between antibodies and cognate receptors. The logic flows that innovation of new capabilities in antibody therapeutics is best served by the discovery of new natural antibody interfaces.

Our characterization of the crystal packing arrangements of 1,456 antibody Fab regions resulted in a diversity of interfaces, with 42 represented in 5 or more PDB entries of nonredundant sequence. The low frequency of most of these together with the generally weak structural and energetic features suggest that many may be crystal artifacts. Confidence in biological relevance is increased by both high prevalence and the existence of shared motifs across multiple results, namely the recurrence of β-sheet dimers and interaction at Fab elbow regions. β-sheet dimers are by far the most commonly observed motif, making up five of the six most prevalent interfaces and present in 30% of nonredundant PDB entries, yet with a diversity of oligomeric and regional architectures. In all cases, the anti-parallel orientations of the paired Fabs, together with their C-terminal distances that exceed hinge flexibility, would preclude intra-Ig Fab interaction, forcing inter-Ig interactions that would promote immune complexation. This notion is supported by the existence of naturally occurring examples where intermolecular β-sheets associate to form protein heterodimers, homodimers, and larger oligomers (Guharoy and Chakrabarti, 2007). Homodimeric β-sheets are observed naturally, for example in ParB (Schumacher and Funnell, 2005), transthyretin (Monaco et al., 1995; Prapunpoj and Leelawatwattana, 2009), and Ras-binding domain of c-Raf1 (Nassar et al., 1995). β-sheet homodimerization has also been successfully used as a template for de novo protein design (Stranges et al., 2011), and is the commonly observed arrangement in the structures of macrocyclic β-sheet peptides (Cheng et al., 2013). The intrinsic preference of exposed β-strands to pair is illustrated most dramatically in the aggregation of amyloid fibrils (Greenwald and Riek, 2010), and it has been proposed that naturally occurring proteins use negative design to avoid edge-to-edge association (Richardson and Richardson, 2002). Indeed a “generic hypothesis” advanced by Dobson and Karplus suggests that extended β-sheets in amyloid structure are an inherent characteristic of polypeptide chains rather than unique to a specific sequence or structure (Dobson and Karplus, 1999). This model is consistent with our observation of intermolecular β-sheet dimers across varying domains and orientations of the antibody Fab. It is tempting to speculate that the Ig fold, in addition to providing loops for diversity and stable scaffolding to support that diversity, also offers antibodies the additional and previously unrealized benefit of avid transient self-association through inter-strand interactions.

Our disulfide engineering experiments provided further support for the discovered interactions and enabled direct structural investigation. Overall, the experimental results suggest that disulfide-trapped species are a result of predisposed contact in solution that, together with favorable geometry, promote coupling of specific homotypic dimers. To explore the functional potential of these interfaces we leveraged a therapeutically relevant antibody-receptor system that is sensitive to oligomeric interaction. The anti-OX40 results represent a first attempt at exploiting this work for optimization, illustrating that the discovered interfaces can serve as novel engineering sites to enhance biotherapeutic properties.

While further biological validation of the discovered interfaces is needed, our results suggest a previously unknown structural feature of antibody Ig domains, and one that would be well-suited for avidity-driven immune response. Transient β-sheet dimers could boost antigen affinity at the BCR level during clonal selection. Transient interfaces could also be relevant at the IgG level for the enhancement of antigen neutralization and Fc-mediated effector functions. The often repetitive and multivalent nature of pathogenic targets, as well as the multiclonal nature of antibody response, have provided evolutionary pressure for the multivalency of isotypes such as IgM and IgA (Kumar et al., 2020; Li et al., 2020) and IgG hexamerization (Diebolder et al., 2014). Such selective pressure has also resulted in more nuanced valency tricks such as chain swap and Fab-dimerization observed in anti-HIV and -SARS-CoV-2 antibodies (Calarese et al., 2003; Williams et al., 2021; Wu et al., 2013), homotypic interactions involved in antibodies against plasmodium (Kucharska et al., 2020), as well as in the context of therapeutic antibodies (Rougé et al., 2020; Tamada et al., 2015). While inter-IgG β-sheet interactions are weak, they could become relevant energetic drivers on the cell surface or in the context of a solution immune complex where the effective concentration of IgG may be high. In this manner, the environment of a cell surface or solution immune complex may be akin to a biomolecular condensate (Feng et al., 2019). While condensates have typically been characterized in the context of intracellular biology, extracellular examples are known, for example contributing to protein assembly in the extracellular matrix and cell-cell adhesion (Chiu et al., 2020; Reichheld et al., 2017). From this perspective, the Fab crystal lattice may be a proxy for how antibodies behave in their condensed native biological environments. In this light the results here, derived from holistic analysis of antibody structural packing data, provide fresh mechanistic insight into how condensation of immune complexes, either at the cell surface or in solution, may enable amplification of immune interactions into immunological response.

## Materials and Methods

### Dataset from SAbDab

All Fab structure coordinates and the corresponding meta-information were downloaded from the SAbDab database (Dunbar et al., 2014) on 01/26/2018. Species and germline information was cross-checked with information extracted from IMGT/3Dstructure-DB (http://www.imgt.org/3Dstructure-DB/) (Ehrenmann et al., 2009; Kaas et al., 2004). Fab PDB structures were then filtered to ensure first that each Fab structure was complete (e.g. both heavy and LCs were longer than 180 residues), and second that the structure was solved by X-ray diffraction with valid symmetry information (e.g “SMTRY” lines in “REMARK 290” section).

### Computational pipeline for identification of antibody interfaces

An in-house computational pipeline was developed to identify symmetric oligomers and their corresponding interfaces. A universal numbering scheme was used for both variable and constant domains to effectively compare the sequence and structures of symmetric oligomers and interfaces across different Fab structures. Unlike a “bottom-up” strategy (e.g. PISA) where all molecular contacts in the crystal lattice are first extracted and later assembled to complexes, we used a “top-down” strategy that first identifies all existing symmetric oligomers and then tracks their corresponding interfaces. Main components of the computational pipeline are detailed below.

#### Universal numbering scheme of Fabs

To analyze sequences and structures across diverse Fabs, a universal numbering scheme was adopted to renumber all positions in each Fab structure. Such universal numbering assigns structurally equivalent positions in different Fabs with the same number. For the variable domain (VH and VL), this was achieved by using the ANARCI program (Dunbar and Deane, 2016) with the AHo convention (Honegger and Plückthun, 2001), a numbering scheme designed to account for gaps in the variable domain to most accurately reflect structural equivalence. For the Fab constant domain (CH1 and CL), a customized numbering scheme referred to as Universal Constant Numbering (UCN) was created as follows. (1) Germline genes (IGHC, IGKC, and IGLC) representing species that appear in the SAbDab dataset were collected from the IMGT database (http://www.imgt.org/vquest/refseqh.html). They were paired with Fab structures containing the highest sequence similarity. IgE sequences were excluded due to lack of available structures in SAbDab. (2) The seven most conserved β-strands corresponding to the protein core were manually identified based on the multiple sequence alignment and structure alignment (if available) of the germline sequences and their paired structures. Interestingly, structures of these β-strands superimpose well even between CH1 and CL. (3) By using these seven intermittent β-strands as fixed regions and accounting for gaps in-between, the representative germline sequences were manually aligned and numbered. The sequence alignment was stored in .stockholm format. (4) Hidden Markov Model (HMM) profile libraries were compiled for each species and germline gene in the manual sequence alignments using the *hmmbuild* and *hmmpress* commands of the HMMER program. (5) All Fab constant domain sequences in the structure dataset were then searched against these pre-computed HMM profile libraries using the *hmmscan* command of the HMMER program, and were mapped to the corresponding positions in the best hit.

#### Identification of interfaces

The asymmetric unit in each Fab PDB structure was expanded into a crystal lattice block by running a PyMOL script that calls the *symexp* command, which expands around the original asymmetric unit up to 30 Å. For computing efficiency, the expanded crystal lattice block was then trimmed down to at most 20 closest Fab monomer structures around the original asymmetric unit, which was large enough to capture almost all potential symmetric interfaces.

To exhaustively identify all interfaces between Fab monomers in each crystal lattice block, all pairs of Fab monomers were examined for all interfaces above 100 Å^2^ (two-sides) after stripping away water and other small molecules. An undirected graph was then built using Fab monomers as nodes and interfaces as edges. An oligomer here is presented as a connected sub-graph, in which every node has at least 1 edge to other nodes in the same sub-graph. By doing this, the problem of searching for all existing oligomers is transformed to searching for all connected sub-graphs. An in-house merging algorithm was developed to efficiently solve this problem.

#### Rotation angles and axis

A transformation matrix was computed between every possible pair of Fab monomers (regardless of contact or not) by calling *align* and *get_object_matrix* PyMOL commands. The rotation angle and axis were mathematically determined from the rotation matrix *R*using the following equations:

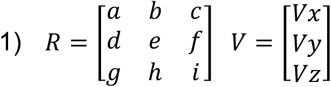

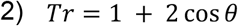

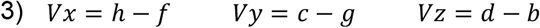

Where *V* is the rotation axis vector, *θ* is the rotation angle, and *Tr* is the trace of *R. θ* is between 0, 180°, which is a sufficient range to effectively test the rotational symmetry. Specially, when *θ* is 180°, to determine the rotation axis, there are the following different possibilities:

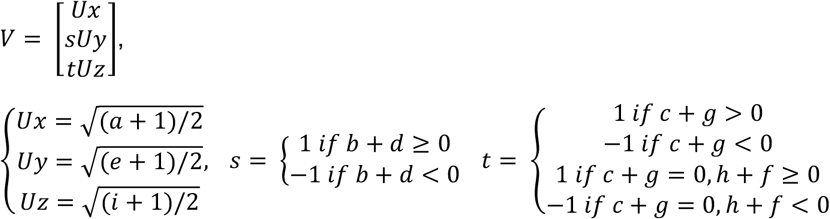

Due to the limited computation precision, imperfect PDB structures, and pseudo symmetry, very similar rotations were treated as identical rotations. To identify them, a pair of rotations were first compared by angles with a cutoff of 5°, and then compared by the angle between their rotation axes with a cutoff of 5°. The angle between rotation axes was calculated using the more numerically stable method:

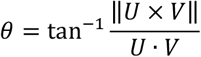

Where *θ* is the angle between axes, and *U* and *V* are the vectors of the two axes. In two special cases, 0° rotations were all dropped, and 180° inter rotation axes (opposite directions) were considered as the same axes.

#### Determination of symmetric oligomers

The last step in the pipeline used the computed rotation information to determine whether a given Fab oligomer is symmetric and in which types. The approach explored every observed rotation in one oligomer. A symmetric rotation allows an oligomer to superimpose to itself. Because a full-atom RSMD calculation on all symmetric rotations for all Fab oligomers was prohibitively expensive, subunits were first represented as centroids that reflected their position and orientation with minimal coordinates. This allowed a fast calculation with a permissive low-resolution RMSD threshold to filter out the vast majority of obvious non-symmetric rotations. A full-atom RMSD calculation was applied to the remaining oligomers. Both centroid and full-atom RMSD were required to be < 5 Å for a given rotation to be considered symmetric. Finally, identical symmetric oligomers were identified and collapsed using the same centroid to full-atom RMSD strategy.

### Clustering analysis

An interface *I* was described using the set of interacting residues between them (identified by using a distance cutoff *R*). Analysis was performed using a distance cutoff of 4 Å and 6 Å.

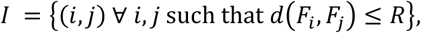

The similarity between two interfaces *I*^*m*^ and *I*^*n*^ is defined as the Jaccard index between the sets of interfaces as follows:

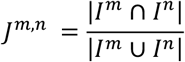

A variant of the analysis utilized a weighted Jaccard similarity, in which each of the two contact pair sets is a real number vector V, with its elements derived from the distance of the corresponding contact pair (and value 0 for the non-contact residue pairs) as follows.

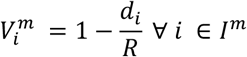

Then, the weighted Jaccard similarity was calculated as

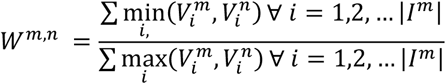

The above similarity values were then used to generate distance matrices (using 1 – J or 1 – W as the values) for distance cutoffs of 4 Å and 6 Å. Hierarchical clustering was then performed using single linkage in R using the ‘hclust’ function. The resulting dendrograms were cut at different heights and the similarities of various cluster members were compared to decide on the optimal cutoff height. The clusters obtained by cutting the Weighted Jaccard index based dendrogram (and interface distance cutoff of 4 Å) at a height of 0.3 was selected for further analysis. Based on the hierarchical clustering, each interface was assigned to an interface cluster.

### Cluster representatives and naming

For each member of a cluster, the average of all weighted Jaccard similarity indices to all other members was calculated. The representative member of each cluster was defined as the interface that had the highest average weighted Jaccard index to all other members.

The name of a given cluster represents the residue at the structural center of the Fab-Fab interface. The name adopts the format X-#, in which X represents commonly used abbreviations for Fab chains (VH, VL, CH1, CL) and # represents the residue number according to either the Kabat or EU numbering convention for variable or constant regions, respectively. The central residue for each cluster was determined as follows. Relevant interface residues for each cluster’s representative member were determined by applying an interface distance cutoff of 5 Å. The xyz coordinates of the α-carbon for each interface residue were averaged to obtain the structural average of the interface. For each residue in the interface, the distance between the given residue’s α-carbon and the structural average xyz coordinates was determined. The residue with the shortest distance to the central xyz coordinates was used for the naming convention. Duplication was avoided in a small number of instances by choosing the second closest residue to the structural center.

### Sequence analysis

For the antibody Fabs in the dataset, all-against-all pairwise alignments were determined for the different Fab regions listed here: VH-only, VL-only, CH1-only, CL-only, and VH/VL-Fv-only. Dynamic programming, with matching residue score at 1, mismatching score at -1, gap-opening score at -6, and gap-extension score at -3, was used to determine the respective alignments. Referencing the pairwise alignment results, Levenshtein distance was calculated to determine the difference between the pairwise-compared regions. Percent-identity was then calculated as the subtraction from 100% by the difference percentage. In the case of Fv percent-identity, the respective VH and VL percent-identities were determined separately; then, the average of the VH- and VL-percent-identities was reported as the identity for the compared Fv regions. In the case of interface percent identity, sequence alignments were performed only at the residue positions of a given interface for each interface (***Supplementary table 2***).

### PISA analysis

Structural features of interfaces were analyzed using the ‘Protein interfaces, surfaces and assemblies’ service PISA at the European Bioinformatics Institute. (http://www.ebi.ac.uk/pdbe/prot_int/pistart.html) (Krissinel and Henrick, 2007). PISA interfaces were downloaded from the PDBePISA site (https://www.ebi.ac.uk/pdbe/pisa/) and mapped to in-house generated oligomer interfaces. Each such oligomer interface consists of single or multiple disconnected PISA interfaces between different chains, and each PISA interface can be included partly or fully, depending on the overlapping interface residues. Buried surface area, number of H-bonds, and number of salt bridges for each interface were calculated as the summation of its component PISA interfaces. In order to accurately analyze the in-house interfaces, we applied two filtering processes for each oligomer interface: (1) removal of PISA interface patches involving antigens, keeping only antibody-antibody interfaces and (2) removal of PISA interfaces for which there was poor overlap (< 0.5) between PISA interface and in-house interface residues.

### Disulfide design

The representative structure of each cluster was used to perform an in silico cysteine scanning simulation using the cysteine scanning module (Salam et al., 2014) in the BioLuminate suite (Schrodinger Inc.).

Default parameters were used, with residues within 5 Å allowed to be flexible (flex_dist = 5.0) while performing stability calculations. Potential disulfide pairs were identified as those residue pairs with a Cβ- Cβ distance within 5.0 Å, irrespective of whether the calculations yielded favorable energies or not. The list of all potential disulfide residue pairs identified through this analysis is summarized in ***Supplementary tables 3-5***.

### DNA construction and protein production

The anti-Her2 and anti-OX40 antibodies used in this work have been described previously (Carter et al., 1992; Yang et al., 2019). Molecular biology to generate the Fab disulfide variants was carried out using gene synthesis (Genewiz). DNAs encoding LCs and Fab HCs in the pRK mammalian expression vector were cotransfected into Expi293 cells for expression. Fabs were purified using CaptureSelect CH1-XL affinity resin (Thermo, 194346201L) followed by SEC using a HiLoad 16/600 Superdex 200 column. Fab monomer and dimer fractions were pooled separately during SEC purification to isolate the desired oligomeric species. Protein quality was assessed by SEC using a Waters xBridge BEH200A SEC 3.5um (7.8 × 300 mm) column (Waters,176003596). For the characterization of oligomeric species following expression, small aliquots of affinity purified Fabs were loaded onto the Waters column, and relative percentages of oligomers were calculated using the area under the chromatogram for each peak. Molecular weight of all Fabs was confirmed by LC/MS. Fabs were stored in a buffer consisting of 20 mM histidine acetate and 150 mM NaCl at pH 5.5.

### Affinity measurements

Solution binding was assessed using a Biacore T200 instrument (GE). Fabs were captured using either a Series S Protein L chip (Cytiva, 29205138) or a human Fab capture reagent immobilized on a Series S CM5 chip (Cytiva, 29104988 and 29234601) according to the manufacturer’s specifications. A 4-fold serial dilution of OX40 starting at 100 nM (G&P Bioscience, FCL0103) was prepared in HBS-P+ buffer (10 mM HEPES, 150 mM NaCl, 0.05% v/v surfactant P20, Cytiva, BR100671) and injected for 5 minutes, followed by a 6 minute dissociation period. Affinity constants were obtained through kinetic fitting using the Biacore Evaluation Software (GE).

### In vitro oxidation

Anti-Her2 Fab disulfide variants were exchanged into a phosphate-buffer saline (PBS) solution at pH 7.4 and concentrated to 0.5 mg/ml using Amicon Ultra centrifugal filters with a 10 kDa molecular weight cutoff (Millipore, UFC901096). Dithiothreitol (Thermo, R0861) was added to 5 mM from a 40 mM stock solution in water. Samples were incubated at 37°C for 2 hours to fully reduce all cysteines. DTT was removed through buffer exchange into PBS using an Amicon Ultra centrifugal filters with a 10 kDa molecular weight cutoff (Millipore, UFC901096). The temperature of the samples was kept at 4°C during the buffer exchange process. A ten-fold molar excess of the oxidizing agent (L)-dehydroascorbic acid (Sigma-Aldrich, 261556) was then added to each sample from a 5 mM stock solution in PBS, and samples were incubated for one week at room temperature. The oxidation reactions were stopped and any remaining free sulfurs were capped upon addition of N-ethylmaleimide (Pierce, 23030) to 5 mM from a 100 mM stock solution in water. SDS-PAGE of the re-oxidized samples was performed on a 4-15% Criterion TGX precast midi protein gel (Bio-Rad, 5671085) under non-reducing conditions. Relative percentages of oligomeric species were calculated using ImageJ.

### Activity assay

The OX40 agonist assay was performed as previously described (Yang et al., 2019). Briefly, OX40 overexpressing Jurkat cells engineered with an NFκB luciferase reporter were seeded at 80,000 cells/well in 20 µl RPMI containing 1% L-glutamine and 10% HI FBS in a 384-well tissue culture plate (Corning Inc., 3985BC). Anti-OX40 antibody formats were serially diluted three-fold in media starting at 30 μg/ml, and 10 µl of the concentrated antibodies were added to each well. For the conditions with crosslinking, 10 µl of AffiniPure goat anti-human IgG Fcγ fragment specific antibody (for IgG samples, Jackson ImmunoResearch Laboratories Inc, 109-005-098) in media was added to yield a 1:1 molar ratio with each antibody dilution. For conditions without crosslinker, 10 µl of media was added to each antibody dilution. The plates were then incubated for 16–18 hrs under 5% CO2 at 37°C. 40 µl of Bright Glo (Promega, E2610) was then added to each well and incubated with shaking at room temperature for 5 min. Luminescence was detected using a Perkin Elmer Envision plate reader. For larger panels of antibodies, automation of this assay was developed using a Tecan Fluent. All activity data from this assay is reported as fold change over control well without antibody.

### Crystallographic structure solution

The purified dimers of the anti-Her2 Fab disulfide variants were concentrated to 10 mg/ml in PBS. Crystallization trials were performed using the sitting-drop vapor diffusion method with commercially available sparse-matrix screens in a 96-well format. The final crystallization condition for the anti-Her2 Fab disulfide variant CH1(S119C) / CH1(G122C) (interface CH1-207) contained 0.8M sodium succinate. The condition for variant VL(G16C) / CL(D170C) (interface VL-108) contained 1% PEG 2000 and 100 mM HEPES pH 8.0. Finally, the condition for variant CL(S202C) / CL(S208C) (interface CL-205) contained 15 % w/v PEG 3350 and 100 mM sodium succinate. Crystals of the dimers were preserved for data collection by brief soaking in a cryo-protectant buffer (25% glycerol added to the reservoir solution), followed by rapid immersion into liquid nitrogen. Diffraction data were collected at the Advanced Light Source (ALS) beam line 5.0.2 for CH1-207 and VL-108, while data for CL-205 were collected at the Northeastern Collaborative Access Team (NE-CAT) beamline 24IDC.

Data were reduced with Global Phasing’s autoProc (Vonrhein et al., 2011) using XDS (Kabsch, 2010). Datasets for CH1-207 and VL-108 were defined with elliptical anisotropic resolution as implemented in the STARANISO procedure (Tickle et al., 2018). Structure determination was done by molecular replacement with Phaser (McCoy et al., 2007) using a search model from a prior internal antibody Fab structure separated into VH, VL, CH1, and CL subdomains. Electron density maps were consistent with the presence of the engineered disulfide bonds at the expected sequence positions (***Supplementary figure 5***) and these were introduced during manual model building in Coot (Emsley and Cowtan, 2004). Of note, the initial solution for VL-108 was determined in a 6-fold symmetry group, but presented only 1 molecule per asymmetric unit in that context with the neighboring linked protomer positioned by apparent crystal symmetry. However, the inability to adequately model the disulfide bond of interest across the special symmetry position and the potential twinning led us to reduce the space group symmetry to P3121 with two Fab molecules per asymmetric unit and to model twinning in the data that had presented as the higher symmetry. Similarly, the CL-205 dataset initially appeared to be in space group P21 with two Fabs per asymmetric unit but had strong indicators of translational pseudo-symmetry (Phenix xtriage (Adams et al., 2010) analysis) and the Rfree value did not decrease much below 30% in refinement efforts. The data were re-processed in P1 and the molecular replacement search repeated to identify 4 Fab molecules per asymmetric unit, and refinement in this setting with amplitude-based twin estimates in REFMAC(Murshudov et al., 1997) provided approximately 4% reduction in R factors. The CH1-207 data were mercifully more straightforward. Models were refined with cycles of REFMAC (Murshudov et al., 1997), Phenix (Adams et al., 2010) or Buster (Bricogne et al., 2011) to reasonable statistics (***Supplementary table 7***). CL-205 has 4 Fab molecules per asymmetric unit in space group P1, the engineered interface between the two Fabs, which are assigned chains H (heavy) and L (light) and the second Fab with chains A (heavy) and B (light) and analogously for the neighboring C/D and E/F chain Fabs. The CH1-207 structure also displays 4 Fab molecules/asu, with engineered interfaces between Fabs HL and AB and Fabs CD and EF. Finally, VL-108 contains 2 Fab molecule/asu in a P3121 space group setting, twinned, assigned chains HL and AB. Coordinates and structure factors are deposited with the PDB under accession codes 7T97 (CH1-207), 7T98 (VL-108), and 7T99 (CL-205).

## Acknowledgements

The authors thank Ingrid Kim, Farzam Farahi, Peter Luan, and Nina Kim for technical contributions, and Kiran Mukhyala for thoughtful input. The synchrotron radiation sources for structure determination are supported by the U.S. Department of Energy, Office of Science, Office of Basic Energy Sciences under Contract No. DE-AC02-05CH11231 (Advanced Light Source) and contract no. DE-AC02-06CH11357 (Advanced Photon Source).

