## Supplementary Information for "Novel Antibody Interfaces Revealed Through Structural Mining"

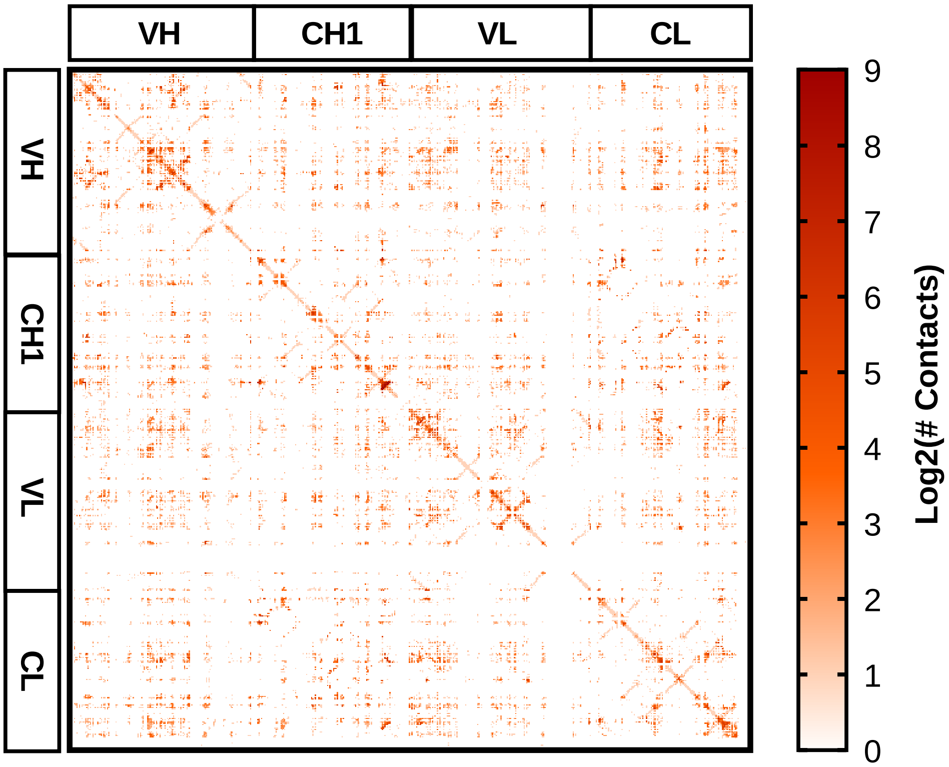

**Supplementary figure 1.** 4 Å contact map of inter-Fab residue pairs. Heavy and light chains are arbitrarily represented by concatenation from left to right and top to bottom. Map coloring on log2 scale reflects the number of times a contact is observed in the lattices of 1456 Fab structures, with 0 being the minimum and 512 being the maximum count.

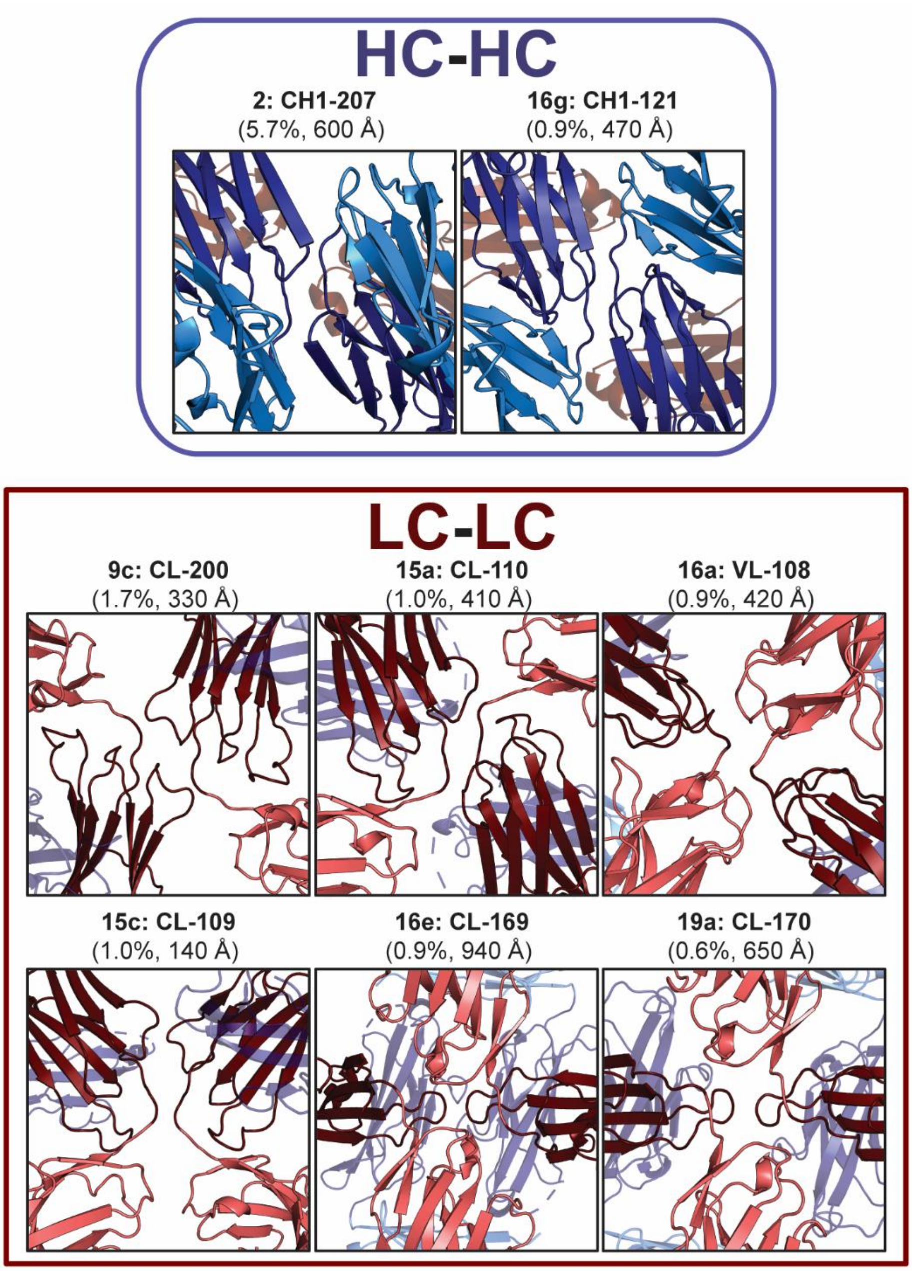

**Supplementary figure 2.** Fab dimers mediated through VH-CH1 and VL-CL elbow regions. Structural similarity between elbow region Fab oligomers that interact as HC-HC (*top*) or LC-LC (*bottom*) dimers. Domains are colored as follows: VH (light blue), CH1 (dark blue), VL (pink), CL (dark red). The labels for each interface include the cluster rank, interface name, nonredundant prevalence percentage, and mean buried surface area.

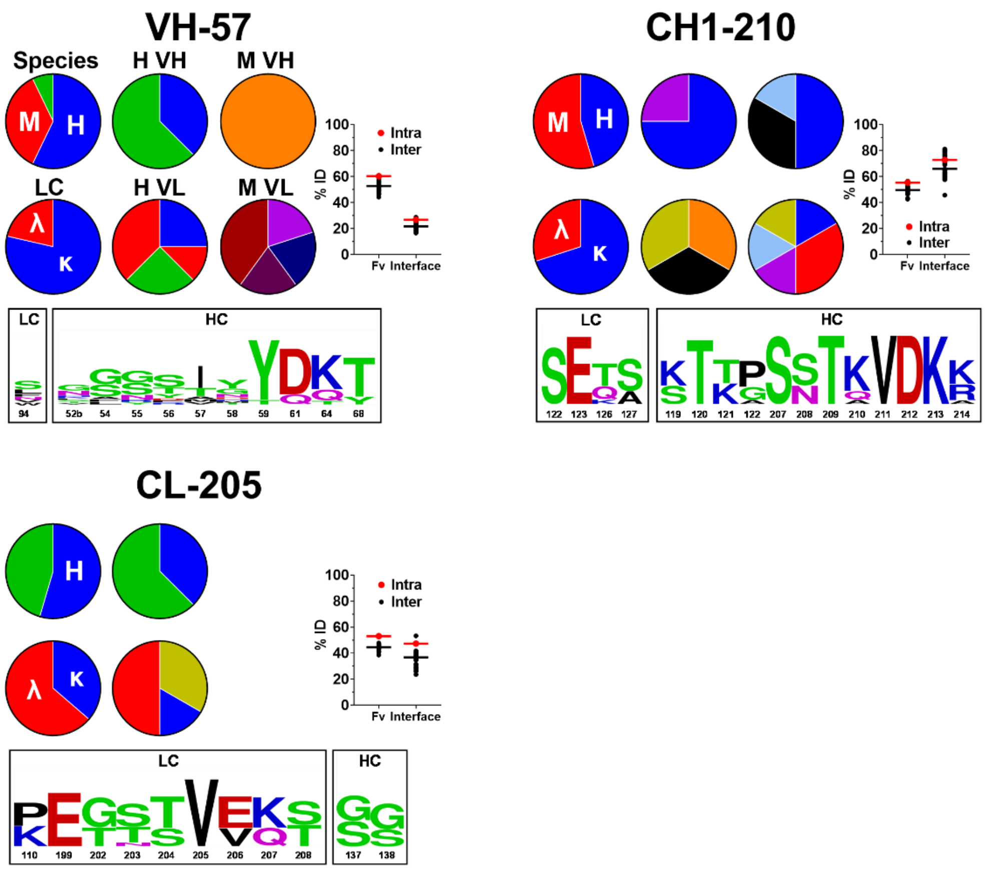

**Supplementary figure 3.** Interface profiles of three β-sheet dimers VH-57, CH1-210, and CL-205. For all interfaces, the upper left-most pie chart provides the sequence distribution of cluster members based on species: human (blue, H), mouse (red, M), and other (e.g., rat, rabbit, etc., green). The lower left-most pie chart provides the sequence distribution of cluster members based on light chain type: kappa (blue, κ), and lambda (red, λ). The four right-most pie charts provide the sequence distribution of human VH (H VH), human VL (H VL), mouse VH (M VH), and mouse VL (M VL) subgroups. The labels above the pie charts in the upper left profile VH-57 are not repeated in the other charts for visual simplicity. The %ID plot on the right provides the intra (red) versus inter (black) cluster sequence identity for both the entire variable region (Fv) as well as only those residues at the interface as shown in the sequence logo and Supplementary Table 2. For the Fv, these values reflect the mean pairwise identities for all VH and VL sequences within the cluster (intra) versus the mean pairwise identities between each member of the cluster and all other members of all other clusters (inter). A similar comparison is made for each interface, where %ID reflects the mean pairwise identity of the residues at the interface of a given cluster aligned with those same residues for each member of the cluster (intra) versus all other members of all other clusters (inter). The sequence logo at the bottom provides weighted sequence composition at interface residues for members of the designated cluster, with numbering according to Kabat and EU conventions for the Fv and constant regions, respectively.

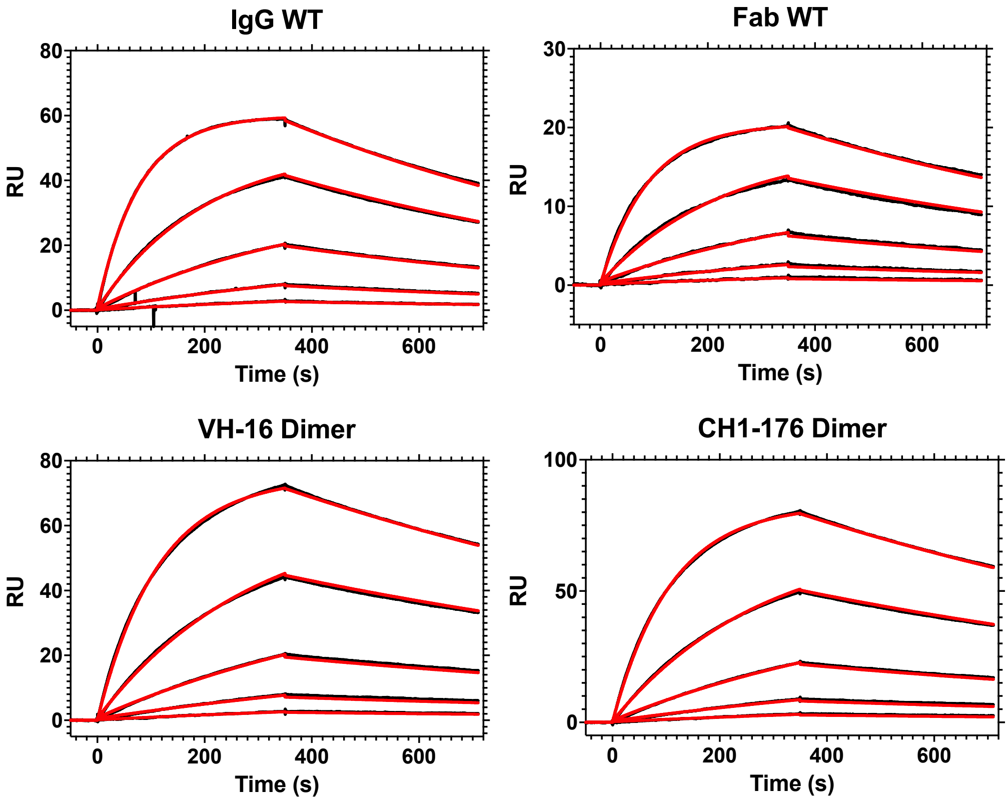

| | Wild-type | | VH-16 | CH1-176 |
|---|---|---|---|---|
| | IgG | Fab | Dimer | |
| **$k_a$ (x$10^5$, $s^{-1}$)** | 1.2 | 1.0 | 0.81 | 0.84 |
| **$k_d$ (x$10^{-3}$, $s^{-1}$ $M^{-1}$)** | 1.2 | 1.1 | 0.78 | 0.83 |
| **$K_D$ (nM)** | 9.9 | 10 | 9.7 | 9.9 |
| **Normalized Analyte Binding*** | 0.40 | 0.48 | 0.52 | 0.57 |

*$R_{max}$ normalized to ligand capture level

**Supplementary figure 4.** OX40 binding by WT IgG and Fab, VH-16 dimer, and CH1-176 dimer by SPR. All antibodies contain the Fv region of the anti-OX40 antibody 3C8. VH-16 represents the Fab dimer of cysteine variant VH(S113C) / CH1(G178C), and CH-176 represents the Fab dimer of cysteine variant VH(P14C) / CL(D151C). Shown are the sensorgrams (*top*) and kinetic fit values and normalized analyte binding (*bottom*). The dimers have slightly slower on/off rates but have the same $K_D$ as the WT Fab and IgG. The normalized binding data are consistent across Fab monomer (WT) and dimer. The value is slightly lower for IgG, but that is expected due to the extra mass of the Fc on the capture ligand.

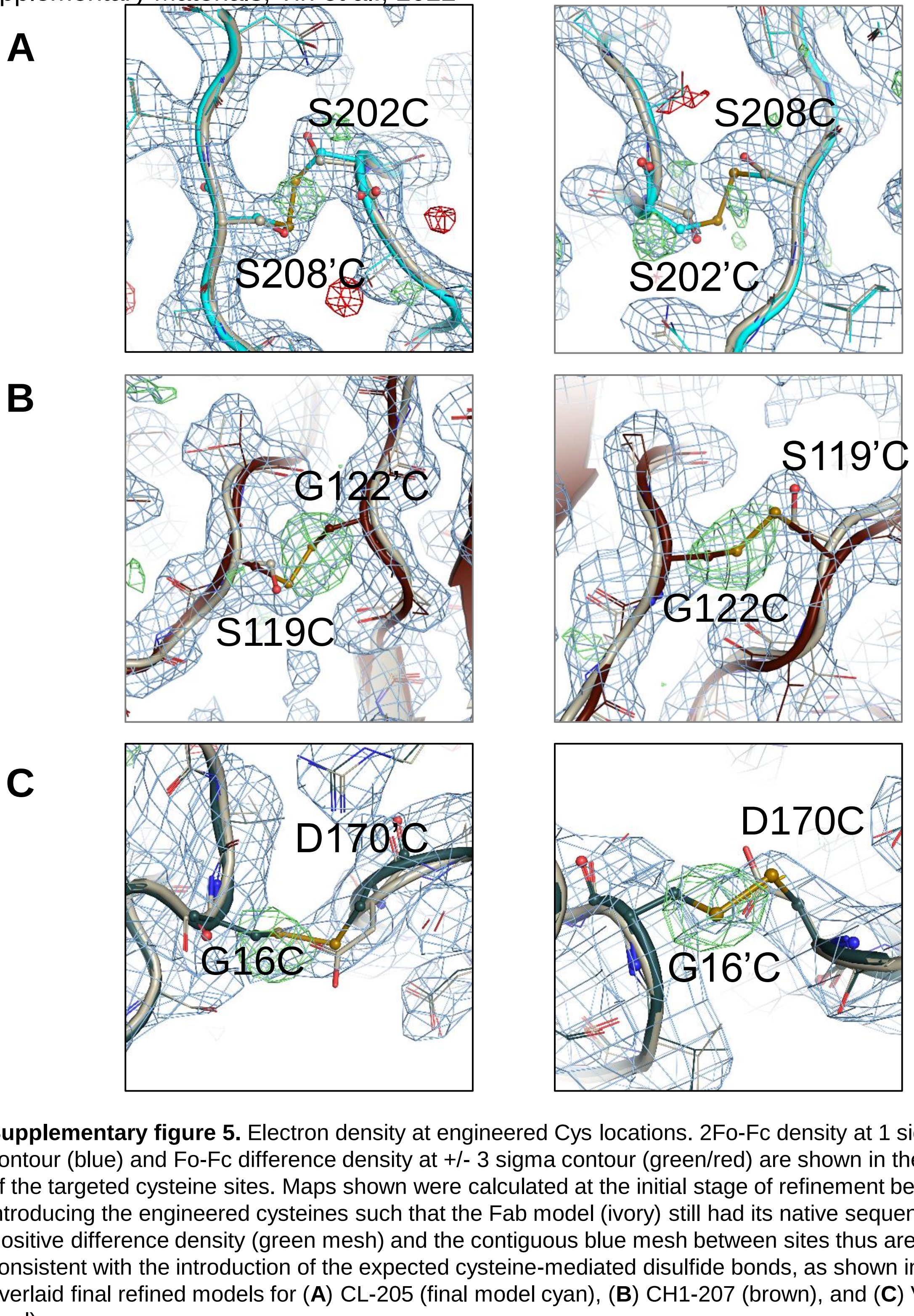

**Supplementary figure 5.** Electron density at engineered Cys locations. 2Fo-Fc density at 1 sigma contour (blue) and Fo-Fc difference density at +/- 3 sigma contour (green/red) are shown in the vicinity of the targeted cysteine sites. Maps shown were calculated at the initial stage of refinement before introducing the engineered cysteines such that the Fab model (ivory) still had its native sequence. Positive difference density (green mesh) and the contiguous blue mesh between sites thus are consistent with the introduction of the expected cysteine-mediated disulfide bonds, as shown in the overlaid final refined models for (**A**) CL-205 (final model cyan), (**B**) CH1-207 (brown), and (**C**) VL-108 (teal).

| Rank | Interface (Kabat, EU) | Total Size (of 1,456) | NR Size (of 981) | NR Prevalence | Area (Å$^2$) | H-Bonds |
|---|---|---|---|---|---|---|
| 1 | CH1-211 | 166 | 133 | 13.6% | 668 ± 134 | 10.4 ± 3.6 |
| 2 | CH1-207 | 61 | 56 | 5.7% | 596 ± 129 | 12.9 ± 3.6 |
| 3 | CH1-209 | 82 | 36 | 3.7% | 397 ± 53 | 6.6 ± 2.6 |
| 4 | CL-211 | 34 | 33 | 3.4% | 678 ± 204 | 10.7 ± 4.7 |
| 5 | VL-11 | 37 | 29 | 3.0% | 768 ± 112 | 14.5 ± 4.3 |
| 6 | CH1-212 | 59 | 27 | 2.8% | 580 ± 220 | 8.9 ± 3.1 |
| 7 | VL-107 | 22 | 19 | 1.9% | 594 ± 58 | 7.5 ± 1.3 |
| 8 | VH-16 | 20 | 18 | 1.8% | 861 ± 291 | 5.9 ± 4.1 |
| 9a | CL-187 | 24 | 17 | 1.7% | 804 ± 180 | 20.5 ± 8 |
| 9b | CH1-176 | 20 | 17 | 1.7% | 646 ± 81 | 12.3 ± 3.5 |
| 9c | CL-200 | 17 | 17 | 1.7% | 335 ± 47 | 6.9 ± 3 |
| 10 | VL-59 | 15 | 15 | 1.5% | 364 ± 126 | 5.9 ± 2.3 |
| 11a | VH-82b | 26 | 14 | 1.4% | 685 ± 159 | 8.8 ± 3.4 |
| 11b | VL-13 | 15 | 14 | 1.4% | 416 ± 92 | 7.2 ± 3.1 |
| 11c | VH-57 | 14 | 14 | 1.4% | 524 ± 140 | 9.2 ± 4.4 |
| 12a | CL-145 | 16 | 13 | 1.3% | 1092 ± 119 | 5.4 ± 3.1 |
| 12b | CH1-161 | 15 | 13 | 1.3% | 1269 ± 145 | 13.4 ± 2.8 |
| 13a | CH1-188 | 13 | 12 | 1.2% | 1397 ± 166 | 12.8 ± 3.2 |
| 13b | VH-82a | 12 | 12 | 1.2% | 306 ± 111 | 8.6 ± 2.3 |
| 14a | CH1-210 | 14 | 11 | 1.1% | 672 ± 149 | 7.4 ± 3.6 |
| 14b | VL-74 | 13 | 11 | 1.1% | 402 ± 27 | 3.2 ± 1.8 |
| 14c | CL-205 | 13 | 11 | 1.1% | 439 ± 89 | 9.7 ± 3.6 |
| 15a | CL-110 | 15 | 10 | 1.0% | 414 ± 79 | 3.8 ± 1.7 |
| 15b | CH1-206 | 13 | 10 | 1.0% | 647 ± 138 | 10.8 ± 3.5 |
| 15c | CL-109 | 10 | 10 | 1.0% | 139 ± 76 | 0.9 ± 0.6 |
| 15d | CH1-197 | 10 | 10 | 1.0% | 421 ± 105 | 4.3 ± 3.3 |
| 16a | VL-108 | 23 | 9 | 0.9% | 417 ± 29 | 5 ± 3.1 |
| 16b | VH-54 | 17 | 9 | 0.9% | 113 ± 59 | 0.2 ± 0.6 |
| 16c | VL-7 | 14 | 9 | 0.9% | 630 ± 197 | 8.8 ± 5 |
| 16d | CH1-139 | 12 | 9 | 0.9% | 556 ± 132 | 8.8 ± 2.3 |
| 16e | CL-169 | 11 | 9 | 0.9% | 945 ± 233 | 9.7 ± 3.6 |
| 16f | CL-151 | 11 | 9 | 0.9% | 227 ± 52 | 4.1 ± 1.5 |
| 16g | CH1-121 | 11 | 9 | 0.9% | 467 ± 42 | 10.5 ± 3.6 |
| 16h | CH1-194 | 10 | 9 | 0.9% | 240 ± 55 | 0.8 ± 1.4 |
| 17a | VL-162 | 15 | 8 | 0.8% | 216 ± 45 | 4.8 ± 0.9 |
| 17b | CH1-165 | 11 | 8 | 0.8% | 1472 ± 168 | 14 ± 3.9 |
| 18a | VL-61 | 11 | 7 | 0.7% | 526 ± 94 | 9.9 ± 2.8 |
| 18b | VH-8 | 10 | 7 | 0.7% | 852 ± 112 | 10.8 ± 2.3 |
| 19a | CL-170 | 10 | 6 | 0.6% | 646 ± 107 | 1.5 ± 1.1 |
| 19b | VL-25 | 10 | 6 | 0.6% | 728 ± 111 | 9.4 ± 4.4 |
| 20a | VH-83 | 12 | 5 | 0.5% | 885 ± 112 | 13.2 ± 2.9 |
| 20b | VH-25 | 10 | 5 | 0.5% | 355 ± 97 | 8.8 ± 5 |

**Supplementary table 1.** Prevalence and structural parameters for the 42 most prevalent interface clusters.

| Rank | Interface | Positions (VL / CL / VH / CH1) |
|---|---|---|
| 1 | CH1-211 | CL(122,123,126) / CH1(119,120,121,122,207,208,209,210,211,212,213,214) |
| 2 | CH1-207 | VH(11,12,13,23,25,72,74,75,112,113) / CH1(118,119,120,206,207,208,209,210) |
| 3 | CH1-209 | CL(126,127) / CH1(119,207,208,209,210,211,212,213) |
| 4 | CL-211 | CL(109,110,112,123,126,202,203,204,205,206,207,208) / CH1(119,120,121,136,137,207,208,209,210,211,212,213,214) |
| 5 | VL-11 | VL(5,6,7,8,9,10,11,12,13,14,17,18,19,20,22,24,70,107) / CL(147,154,155,156) |
| 6 | CH1-212 | CL(151,152,153,154,155,156,157,158) / CH1(160,176,210,211,212,213,214) |
| 7 | VL-107 | VL(14,107) / CL(108,110,111,112,140,141,160,199,200,201,202) / VH(112,113) / CH1(118,119,150,174,175,176,177,178,179,180) |
| 8 | VH-16 | VL(1,3,5,6,7,9,24,26) / CL(160) / VH(13,14,15,16,41,43,84,85,113) / CH1(174,175,176,177,178) |
| 9a | CL-187 | CL(155,157,182,184,185,187,188,189) / CH1(159,162,163,165,188,189,192,193,196,197,198) |
| 9b | CH1-176 | CL(151,152,153,154,155,185,188,189) / VH(14,82c,83,84,85,113) / CH1(118,119,176,178,179) |
| 9c | CL-200 | VL(107) / CL(108,109,110,140,198,199,200,201,202,203) |
| 10 | VL-59 | VL(59,60,61,76,77,79,80,81) |
| 11a | VH-82b | VH(14,15,61,62,63,64,65,66,82a,82b,82c,83,84,85) |
| 11b | VL-13 | VL(8,11,12,13,14,17,18,19,20,107) / CL(199) |
| 11c | VH-57 | VL(94) / VH(52b,54,55,56,57,58,59,61,64,68) |
| 12a | CL-145 | VL(3,4,5,7,8,9,10,11,12,24,26,27,98,99,100) / CL(143,145,149,151,152,190,191,195,197,199,202,203,204,206,208,210) |
| 12b | CH1-161 | CL(138) / VH(1,3,5,7,23,24,25,26,75,76,77,105) / CH1(138,139,157,160,161,162,165,166,187,188,189,191,192,195,196,205,210) |
| 13a | CH1-188 | VH(42,84,85,87,110,111,112) / CH1(132,135,136,137,138,139,150,152,165,171,172,173,174,175,176,180,187,188,189,190,191,192,193,194,195,196) |
| 13b | VH-82a | VH(15,16,17,65,66,68,81,82a,82b,82c) |
| 14a | CH1-210 | CL(122,123,126,127) / CH1(119,120,121,122,207,208,209,210,211,212,213,214) |
| 14b | VL-74 | VL(17,18,20,52,63,64,65,67,74,76) |
| 14c | CL-205 | CL(110,199,202,203,204,205,206,207,208) / CH1(137,138) |

**Supplementary table 2 (Part 1).**

| Rank | Interface | Positions (VL / CL / VH / CH1) |
|---|---|---|
| 15a | CL-110 | VL(107) / CL(108,109,110,112,140,199,200,201,202,203) |
| 15b | CH1-206 | VH(3,11,12,13,16,24,25,74,75,76,112,113) / CH1(118,119,120,205,206,207,208,209,210) |
| 15c | CL-109 | CL(108,109,110) |
| 15d | CH1-197 | CL(157,159,160) / VH(84,85) / CH1(160,161,174,175,176,194,195,197) |
| 16a | VL-108 | VL(14,15,16,17,107) / CL(108,109,169,170,172) |
| 16b | VH-54 | VH(52b,54) |
| 16c | VL-7 | VL(5,6,7,8,9,10,22,24,67,70) / CL(143) |
| 16d | CH1-139 | CL(108,109,114,138,169,170,171,172) / CH1(137,138,139,191,192,193,195) |
| 16e | CL-169 | VL(15,61,79,80,81) / CL(108,109,168,169,170,171) / VH(1,3,4,5,105) / CH1(137,155,157,160,161,162,189,190,191) |
| 16f | CL-151 | CL(149,151,152,188,189,190,191) |
| 16g | CH1-121 | VH(13) / CH1(119,120,121,122,123,124,205,206,207,208,209,210,211,212) |
| 16h | CH1-194 | CH1(194,195,197,217,218,219) |
| 17a | VL-162 | VL(60,61) / CH1(161,162,163,164,165) |
| 17b | CH1-165 | CL(108,170) / VH(1,3,5,7,21,23,24,25,75,76,105) / CH1(138,139,155,157,160,161,162,165,189,191,192,203,205,210,212) |
| 18a | VL-61 | VL(15,18,52,54,55,56,57,58,59,60,61,62,63,64,65,74,76,77,79) |
| 18b | VH-8 | VH(7,8,9,10,13,16,17,19,21,30,52a,54,71,72,73,75,77,79,81,82a,107,108) / CH1(205,206,208) |
| 19a | CL-170 | VL(15,16,56,57,58,59,60,80) / CL(108,109,168,169,170) / CH1(138,189,191) |
| 19b | VL-25 | VL(1,3,25,26,27,67,68,69,70) / CL(144,145,159,160,161,163) / CH1(173,174,175,176) |
| 20a | VH-83 | CL(154,155,156,157,158,160,181,188) / VH(14,15,61,62,64,65,66,68,82b,82c,83,113) / CH1(174,175,176,177) |
| 20b | VH-25 | VH(1,25,26,27,28,29,73,74,76) |

**Supplementary table 2 (Part 2).** Residue positions of the 42 most prevalent interfaces. Positions are numbered according to Kabat for the VL/VH and EU for the CL/CH1.

| Rank | Interface | Variant | % Monomer | % Dimer | % Higher Order |
|---|---|---|---|---|---|
| 1 | CH1-211 | CH1(N208C) / CH1(K214C) | 74 | 24 | |
| 1 | CH1-211 | CH1(K210C) / CH1(D212C) | 59 | 39 | |
| 2 | CH1-207 | VH(L11C) / CH1(N208C) | 84 | 16 | |
| 2 | CH1-207 | CH1(S119C) / CH1(G122C) | 53 | 47 | |
| 2 | CH1-207 | CH1(S119C) / CH1(S207C) | 96 | 4 | |
| 2 | CH1-207 | CH1(S119C) / CH1(T209C) | 70 | 29 | |
| 2 | CH1-207 | CH1(T120C) | 99 | 1 | |
| 2 | CH1-207 | CH1(T120C) / CH1(S207C) | 98 | 2 | |
| 3 | CH1-209 | CH1(N208C) / CH1(D212C) | 60 | 39 | |
| 3 | CH1-209 | CH1(K210C) | 99 | | |
| 4 | CL-211 | CH1(S119C) / CH1(S190C) | 80 | 19 | |
| 8 | VH-16 | VH(P14C) / CH1(L174C) | 90 | 10 | |
| 8 | VH-16 | VH(S113C) / CH1(G178C) | 62 | 25 | 13 |
| 8 | VH-16 | VH(Q13C) / CH1(S176C) | 83 | 16 | |
| 10 | VL-59 | VH(E1C) | 95 | | 4 |
| 11a | VH-82b | VH(P14C) / VH(K64C) | 50 | 47 | 3 |
| 11a | VH-82b | VH(P14C) / VH(G65C) | 39 | 60 | |
| 11a | VH-82b | VH(D61C) / VH(A84C) | 70 | 30 | |
| 11a | VH-82b | VH(N82aC) / VH(S82bC) | 67 | 31 | |
| 11a | VH-82b | VH(S82bC) | 67 | 28 | |
| 11a | VH-82b | VH(G65C) / VH(R83C) | 49 | 50 | |
| 11a | VH-82b | VH(G15C) / VH(G65C) | 9 | 84 | |
| 11c | VH-57 | VH(T57C) | 99 | | |
| 11c | VH-57 | VH(N54C) / VH(K64C) | 28 | 32 | 38 |
| 12b | CH1-161 | VH(V5C) / CH1(S192C) | 68 | 22 | 9 |
| 12b | CH1-161 | VH(S7C) / CH1(T195C) | 79 | 19 | |
| 12b | CH1-161 | VH(A23C) / CH1(P189C) | 90 | 10 | |
| 12b | CH1-161 | VH(A23C) / CH1(S191C) | 95 | 5 | |
| 12b | CH1-161 | VH(A23C) / CH1(S192C) | 66 | 32 | |
| 12b | CH1-161 | VH(S25C) / CH1(T139C) | 89 | 11 | |
| 12b | CH1-161 | VH(S25C) / CH1(P189C) | 88 | 11 | |
| 12b | CH1-161 | VH(S25C) / CH1(G138C) | 80 | 10 | 9 |
| 12b | CH1-161 | VH(K75C) / CH1(S191C) | 91 | 9 | |
| 12b | CH1-161 | CH1(T155C) / CH1(A162C) | 90 | 10 | |
| 12b | CH1-161 | CH1(S157C) / CH1(G161C) | 88 | 12 | |
| 12b | CH1-161 | CH1(S160C) / CH1(N201C) | 73 | 23 | 4 |
| 12b | CH1-161 | VH(T77C) / CH1(S191C) | 97 | 3 | |
| 12b | CH1-161 | CH1(G161C) | 93 | 7 | |

**Supplementary table 3 (Part 1).**

| Rank | Interface | Variant | % Monomer | % Dimer | % Higher Order |
|---|---|---|---|---|---|
| 13a | CH1-188 | VH(P41C) / CH1(S190C) | 38 | 45 | 17 |
| 13a | CH1-188 | VH(P41C) / CH1(S191C) | 41 | 56 | 3 |
| 13a | CH1-188 | VH(G42C) / CH1(L193C) | 78 | | 21 |
| 13a | CH1-188 | VH(T110C) / CH1(G138C) | 89 | | 11 |
| 13a | CH1-188 | VH(S112C) / CH1(G137C) | 82 | 9 | 8 |
| 13a | CH1-188 | CH1(L174C) / CH1(P189C) | 95 | 5 | |
| 13a | CH1-188 | CH1(S165C) / CH1(S176C) | 45 | 55 | |
| 13b | VH-82a | VH(G15C) | 86 | 14 | |
| 13b | VH-82a | VH(S17C) / VH(S82bC) | 90 | 10 | |
| 13b | VH-82a | VH(R66C) / VH(N82aC) | 85 | 14 | |
| 15d | CH1-197 | VH(E85C) / CH1(G161C) | 99 | | |
| 16d | CH1-139 | CH1(G138C) / CH1(T139C) | 93 | 7 | |
| 16d | CH1-139 | CH1(T139C) | 94 | 6 | |
| 16e | CL-169 | VH(Q3C) / CH1(S191C) | 90 | 10 | |
| 16e | CL-169 | CH1(S160C) | 96 | 4 | |
| 16g | CH1-121 | CH1(T120C) / CH1(G122C) | 80 | 19 | |
| 16g | CH1-121 | CH1(K121C) | 99 | | |
| 16h | CH1-194 | CH1(G194C) | 95 | 5 | |
| 16h | CH1-194 | CH1(G194C) / CH1(T195C) | 92 | 8 | |
| 17a | CH1-162 | VH(G99C) / CH1(T195C) | 65 | 32 | |
| 17b | CH1-165 | VH(A23C) / CH1(G138C) | 84 | 11 | 4 |
| 17b | CH1-165 | VH(N76C) / CH1(G138C) | 87 | 12 | |
| 17b | CH1-165 | VH(Q105C) / CH1(S165C) | 99 | 1 | |
| 17b | CH1-165 | CH1(S160C) / CH1(N203C) | 92 | 7 | |
| 17b | CH1-165 | VH(S7C) / CH1(P189C) | 93 | 7 | |
| 17b | CH1-165 | CH1(A162C) / CH1(N203C) | 98 | 2 | |
| 18b | VH-8 | VH(G9C) / VH(Y79C) | 91 | 9 | |
| 18b | VH-8 | VH(G10C) / VH(D72C) | 93 | 7 | |
| 18b | VH-8 | VH(S17C) | 82 | 18 | |
| 18b | VH-8 | VH(R19C) | 98 | 2 | |
| 18b | VH-8 | VH(T73C) / CH1(K205C) | 97 | 3 | |
| 20a | VH-83 | VH(G15C) / CH1(S176C) | 16 | 51 | 29 |
| 20a | VH-83 | VH(G15C) / CH1(S177C) | 27 | 67 | 6 |
| 20a | VH-83 | VH(S82bC) / CH1(S176C) | 34 | 45 | 18 |
| 20a | VH-83 | VH(R83C) / CH1(S176C) | 89 | 11 | |
| 20a | VH-83 | VH(S113C) | 67 | 29 | 4 |
| 20b | VH-25 | VH(S25C) | 97 | 2 | |
| 20b | VH-25 | VH(S25C) / VH(N76C) | 84 | 15 | |
| 20b | VH-25 | VH(G26C) / VH(N76C) | 96 | 4 | |
| WT | WT | WT | 97 | 3 | |
| WT | WT | WT | 99 | 1 | |

**Supplementary table 3 (Part 2).** Cysteine variants at heavy chain / heavy chain (HCHC) interfaces. Data represent normalized integrated peaks of monomer, dimer, and higher order (> dimer) species from analytical SEC. Due to the nature of the peak integration the results do not necessarily add to precisely 100%. Variants are labeled according to Kabat for the VH and EU for the CH1.

| Rank | Interface | Variant | % Monomer | % Dimer | % Higher Order |
|---|---|---|---|---|---|
| 4 | CL-211 | CL(T109C) / CL(K126C) | 90 | 3 | 6 |
| 5 | VL-11 | VL(S7C) / VL(D17C) | 67 | 24 | 10 |
| 5 | VL-11 | VL(S7C) / VL(R18C) | 66 | 26 | |
| 5 | VL-11 | VL(P8C) / VL(L11C) | 89 | 11 | |
| 5 | VL-11 | VL(P8C) / VL(A13C) | 82 | 11 | |
| 5 | VL-11 | VL(S9C) / VL(K107C) | 83 | 7 | 8 |
| 5 | VL-11 | VL(S10C) / VL(S12C) | 76 | 22 | 12 |
| 5 | VL-11 | VL(L11C) | 87 | 12 | |
| 7 | VL-107 | VL(P8C) / CL(S156C) | 58 | 34 | 5 |
| 7 | VL-107 | VL(P8C) / CL(G157C) | 84 | 8 | 8 |
| 9c | CL-200 | CL(V110C) / CL(S202C) | 75 | 19 | 4 |
| 11b | VL-13 | VL(R18C) | 87 | 11 | |
| 12a | CL-145 | VL(Q3C) / CL(K190C) | 84 | 7 | 8 |
| 12a | CL-145 | VL(Q3C) / CL(V191C) | 89 | 10 | |
| 12a | CL-145 | VL(S7C) / CL(T206C) | 72 | 21 | 4 |
| 12a | CL-145 | VL(P8C) / CL(S203C) | 85 | 3 | 11 |
| 12a | CL-145 | VL(S26C) / CL(K190C) | 83 | 8 | 7 |
| 12a | CL-145 | VL(Q100C) / CL(N152C) | 88 | 2 | 8 |
| 12a | CL-145 | CL(Q199C) | 88 | 11 | |
| 12a | CL-145 | VL(S9C) / CL(P204C) | 88 | 11 | |
| 14a | CH1-210 | CL(S127C) | 88 | 11 | |
| 14a | CH1-210 | CL(G128C) | 88 | 11 | |
| 14b | VL-74 | VL(R18C) / VL(S63C) | 84 | 10 | |
| 14b | VL-74 | VL(R18C) / VL(S65C) | 80 | 18 | |
| 14c | CL-205 | CL(S202C) / CL(S208C) | 62 | 28 | 4 |
| 14c | CL-205 | CL(P204C) / CL(T206C) | 86 | 4 | 9 |
| 15a | CL-110 | CL(T109C) / CL(A112C) | 57 | 31 | 6 |
| 15a | CL-110 | CL(T109C) / CL(G200C) | 83 | 11 | 5 |
| 15a | CL-110 | CL(T109C) / CL(L201C) | 78 | 12 | 7 |
| 15a | CL-110 | CL(V110C) / CL(G200C) | 87 | 12 | |

**Supplementary table 4 (Part 1).**

| Rank | Interface | Variant | % Monomer | % Dimer | % Higher Order |
|---|---|---|---|---|---|
| 15c | CL-109 | CL(T109C) | 89 | 11 | |
| 16a | VL-108 | VL(G16C) / CL(K169C) | 54 | 38 | 7 |
| 16a | VL-108 | VL(G16C) / CL(D170C) | 45 | 50 | 5 |
| 16c | VL-7 | VL(S7C) | 78 | 18 | 3 |
| 16c | VL-7 | VL(S7C) / VL(R24C) | 79 | 17 | |
| 16c | VL-7 | VL(P8C) / VL(D70C) | 89 | 10 | |
| 16c | VL-7 | VL(S67C) / CL(E143C) | 85 | 14 | |
| 16c | VL-7 | VL(G68C) / CL(E143C) | 75 | 18 | 5 |
| 16e | CL-169 | VL(P80C) / CL(D170C) | 89 | 10 | |
| 16e | CL-169 | CL(S168C) / CL(K169C) | 92 | 8 | |
| 17b | CH1-165 | CL(D151C) | 54 | 41 | 5 |
| 17b | CH1-165 | CL(D151C) / CL(K190C) | 78 | 16 | 4 |
| 17b | CH1-165 | CL(D151C) / CL(V191C) | 81 | 17 | |
| 17b | CH1-165 | CL(N152C) | 88 | 10 | |
| 18a | VL-61 | VL(R18C) / VL(S52C) | 81 | 17 | |
| 18a | VL-61 | VL(S60C) | 79 | 15 | 5 |
| 18a | VL-61 | VL(S60C) / VL(R61C) | 89 | 11 | |
| 18a | VL-61 | VL(S63C) / VL(S76C) | 87 | 12 | |
| 19a | CL-170 | VL(P80C) / CL(K169C) | 90 | 10 | |
| 19a | CL-170 | VL(G16C) / CL(T109C) | 52 | 37 | 7 |
| 19b | VL-25 | VL(Q3C) / CL(S156C) | 66 | 22 | 10 |
| 19b | VL-25 | VL(T5C) / CL(G157C) | 73 | 5 | 21 |
| 19b | VL-25 | VL(S26C) / CL(S159C) | 88 | 11 | |
| 19b | VL-25 | VL(Q27C) / CL(E161C) | 88 | 12 | |
| WT | WT | WT | 99 | | |
| WT | WT | WT | 99 | | |

**Supplementary table 4 (Part 2).** Cysteine variants at light chain / light chain (LCLC) interfaces. Data represent normalized integrated peaks of monomer, dimer, and higher order (> dimer) species from analytical SEC. Due to the nature of the peak integration the results do not necessarily add to precisely 100%. Variants are labeled according to Kabat for the VL and EU for the CL.

| Rank | Interface | Variant | % Monomer | % Dimer | % Higher Order |
|---|---|---|---|---|---|
| 3 | CH1-209 | CH1(S119C) / CL(K126C) | 62 | 36 | |
| 4 | CL-211 | CH1(K213C) / CL(S203C) | 89 | 7 | |
| 4 | CL-211 | CH1(D212C) / CL(P204C) | 97 | | |
| 4 | CL-211 | CH1(K210C) / CL(T206C) | 96 | | |
| 4 | CL-211 | CH1(N208C) / CL(S208C) | 85 | 14 | |
| 6 | CH1-212 | CH1(K210C) / CL(A153C) | 86 | 11 | |
| 6 | CH1-212 | CH1(D212C) / CL(A153C) | 87 | 12 | |
| 6 | CH1-212 | CH1(K213C) / CL(S156C) | 66 | 29 | |
| 6 | CH1-212 | CH1(K214C) / CL(N158C) | 91 | 5 | |
| 6 | CH1-212 | CH1(S160C) / CL(K188C) | 90 | 9 | |
| 6 | CH1-212 | CH1(T197C) / CL(S182C) | 94 | 4 | 2 |
| 6 | CH1-212 | CH1(K214C) / CL(G157C) | 90 | 4 | 5 |
| 7 | VL-107 | CH1(L174C) / VL(S14C) | 94 | | |
| 7 | VL-107 | CH1(L174C) / VL(K107C) | 95 | | |
| 7 | VL-107 | CH1(A118C) / CL(V110C) | 92 | | |
| 7 | VL-107 | CH1(G178C) / CL(V110C) | 95 | | |
| 7 | VL-107 | VH(S113C) / CL(A112C) | 67 | 29 | |
| 7 | VL-107 | CH1(A118C) / CL(A112C) | 92 | | |
| 7 | VL-107 | CH1(G178C) / CL(P141C) | n.a. | | |
| 7 | VL-107 | CH1(S119C) / CL(S202C) | 72 | 19 | |
| 7 | VL-107 | CH1(A118C) / CL(G200C) | 92 | | |
| 9a | CL-187 | CH1(A162C) / CL(S182C) | 91 | | |
| 9b | CH1-176 | VH(P14C) / CL(D151C) | 65 | 33 | |
| 9b | CH1-176 | VH(R83C) / CL(A153C) | 85 | 7 | 7 |
| 9b | CH1-176 | VH(R83C) / CL(N152C) | 87 | 6 | 6 |
| 9b | CH1-176 | VH(A84C) / CL(A153C) | 89 | 4 | 6 |

**Supplementary table 5 (Part 1).**

| Rank | Interface | Variant | % Monomer | % Dimer | % Higher Order |
|---|---|---|---|---|---|
| 13a | CH1-188 | CH1(T164C) / CL(G157C) | 74 | 11 | 14 |
| 13a | CH1-188 | CH1(T195C) / CL(E165C) | 91 | | |
| 14c | CL-205 | CH1(T135C) / CL(V110C) | 94 | | |
| 15d | CH1-197 | CH1(L193C) / CL(G157C) | 92 | | |
| 15d | CH1-197 | CH1(G194C) / CL(G157C) | 88 | 5 | 6 |
| 16d | CH1-139 | CH1(G138C) / CL(S114C) | 89 | 11 | |
| 16d | CH1-139 | CH1(S191C) / CL(A111C) | 94 | | |
| 16d | CH1-139 | CH1(S191C) / CL(N138C) | 95 | | |
| 16d | CH1-139 | CH1(S191C) / CL(D170C) | 61 | 39 | |
| 16d | CH1-139 | CH1(S192C) / CL(D170C) | 61 | 36 | |
| 16d | CH1-139 | CH1(G194C) / CL(T109C) | 95 | | |
| 16d | CH1-139 | CH1(T195C) / CL(D170C) | 76 | 21 | |
| 16d | CH1-139 | CH1(G137C) / CL(S114C) | 97 | | |
| 16d | CH1-139 | CH1(S191C) / CL(T172C) | 97 | | |
| 17a | CH1-162 | CH1(A162C) / VL(P59C) | 92 | | |
| 17b | CH1-165 | VH(E1C) / CL(A111C) | 87 | 5 | 7 |
| 19a | CL-170 | CH1(G138C) / VL(P59C) | 96 | | |
| 19b | VL-25 | CH1(V173C) / VL(D28C) | 95 | | |
| 19b | VL-25 | CH1(L174C) / VL(G68C) | 92 | | |
| 19b | VL-25 | CH1(Q175C) / VL(T69C) | 97 | | |
| 20a | VH-83 | VH(G65C) / CL(Q155C) | 96 | | |
| 20a | VH-83 | VH(G65C) / CL(N158C) | 90 | 9 | |
| 20a | VH-83 | VH(R66C) / CL(G157C) | 91 | | |
| WT | WT | WT | 99 | | |

**Supplementary table 5 (Part 2).** Cysteine variants at heavy chain / light chain (HCLC) interfaces. Data represent normalized integrated peaks of monomer, dimer, and higher order (> dimer) species from analytical SEC. Due to the nature of the peak integration the results do not necessarily add to precisely 100%. Variants are labeled according to Kabat for the VH/VL and EU for the CH1/CL.

| Rank | Interface | HC Variant | LC Variant | Expression (%) | | | In vitro oxidation (%) | | |
|---|---|---|---|---|---|---|---|---|---|
| | | | | Mon | Dim | HO | Mon | Dim | HO |
| 2 | CH1-207 | CH1(S119C) / CH1(G122C) | | 53 | 47 | | 65 | 35 | |
| 11a | VH-82b | VH(G15C) / VH(G65C) | | 9 | 84 | | 34 | 50 | 17 |
| 12b | CH1-161 | VH(A23C) / CH1(S192C) | | 66 | 32 | | 70 | 29 | 1 |
| 20a | VH-83 | VH(G15C) / CH1(S176C) | | 16 | 51 | 29 | 35 | 63 | 2 |
| 5 | VL-11 | | VL(S7C) / VL(D17C) | 67 | 24 | 10 | 59 | 41 | |
| 14c | CL-205 | | CL(S202C) / CL(S208C) | 62 | 28 | 4 | 45 | 52 | 4 |
| 15a | CL-110 | | CL(T109C) / CL(A112C) | 57 | 31 | 6 | 52 | 48 | |
| 16a | VL-108 | | VL(G16C) / CL(D170C) | 45 | 50 | 5 | 52 | 48 | |
| 19a | CL-170 | | VL(G16C) / CL(T109C) | 52 | 37 | 7 | 43 | 54 | 3 |
| 9b | CH1-176 | VH(P14C) | CL(D151C) | 65 | 33 | | 52 | 37 | 11 |
| 16d | CH1-139 | CH1(S191C) | CL(D170C) | 61 | 39 | | 24 | 68 | 8 |
| | MM1 | VH(P14C) | CL(D170C) | | | | 51 | 32 | 17 |
| | MM2 | CH1(S191C) | CL(D151C) | | | | 8 | 26 | 66 |
| | MM3 | CH1(S119C) | CL(D170C) | | | | 44 | 32 | 24 |
| | MM4 | CH1(S191C) | CL(K126C) | | | | 30 | 34 | 37 |

**Supplementary table 6.** In vitro assembly of select cysteine variants. Data represent normalized integrated peaks of monomer (Mon), dimer (Dim), and higher order (HO, > dimer) species from both expression and in vitro oxidation. Due to the nature of the peak integration the results do not necessarily add to precisely 100%. Control mismatched (MM) variants represent in vitro assembled heavy and light chains cysteine variants that were not designed based on the discovered interfaces. Variants are labeled according to Kabat for the VH/VL and EU for the CH1/CL.

| | CH1-207 (7T97) | VL-108 (7T98) | CL-205 (7T99) |
|---|---|---|---|
| Wavelength | 0.9795 | 0.9999 | 0.9792 |
| Resolution range | 75.69 - 2.144 (2.297 - 2.144) | 67.01 - 2.906 (3.01 - 2.906) | 61.35 - 2.651 (2.746 - 2651) |
| Space group | C 1 2 1 | P 31 2 1 | P 1 |
| Unit cell | 151.377 141.075 118.605<br>90 90.323 90 | 99.634 99.634 212.675<br>90 90 120 | 52.824 62.451 124.018<br>89.992 98.374 89.893 |
| Anisotropic Data Analysis | STARANISO | STARANISO | N/A |
| Diffraction limits | 2.998 2.146 2.144 | 3.548 3.548 2.817 | - |
| Eigenvector-1 | 0.999 0.000 -0.040 | 1 0 0 | - |
| Eigenvector-2 | 0.000 1.000 0.000 | 0 1 0 | - |
| Eigenvector-3 | 0.040 0.000 0.999 | 0 0 1 | - |
| Direction-1 | 0.999 _a_*-0.036 _c_* | 0.894 _a_*-0.447 _b_* | - |
| Direction-2 | _b_* | _b_* | - |
| Direction-3 | 0.052 _a_*+0.999 _c_* | _c_* | - |
| Total reflections | 313557 (16196) | 177656 (8545) | 69121 (7639) |
| Unique reflections | 90662 (4534) | 18037 (903) | 37752 (4287) |
| Multiplicity | 3.5 (3.6) | 9.8 (9.5) | 1.8 (31.9.5) |
| Completeness (spherical, %) | 66.59 (17.9) | 69.3 (16.0) | 95.55 (9424) |
| Completeness (ellipsoidal, %) | 92.5 (64.8) | 93.2 (69.5) | - |
| Mean I/sigma(I) | 7.8 (1.5) | 7.9 (1.6) | 15.45 (4.68) |
| Wilson B-factor | 32.54 | 65.36 | 62.95 |
| R-merge | 0.126 (0.921) | 0.319 (1.46) | 0.02947 (0.1667) |
| R-meas | 0.149 (1.085) | 0.337 (1.544) | 0.04165 (0.2357) |
| R-pim | 0.080 (0.570) | 0.107 (0.498) | 0.02942 (0.1665) |
| CC1/2 | 0.995 (0.502) | 0.992 (0.609) | 0.999 (0.974) |
| CC* | 0.998 (0.614) | 0.997 (0.647) | 1 (0.993) |
| Reflections used in refinement | 90617 (1731) | 18568 (166) | 43501 (4285) |
| Reflections used for R-free | 4655 (110) | 923 (6) | 1668 (164) |
| R-work | 0.2057 (0.2914) | 0.1955 (0.1878) | 0.2114 (0.2247) |
| R-free | 0.2370 (0.3377) | 0.2463 (0.1192) | 0.2797 (0.3476) |
| CC(work) | 0.932 (0.652) | 0.840 (0.613) | 0.924 (0.658) |
| CC(free) | 0.902 (0.541) | 0.781 (0.888) | 0.937 (0.639) |
| Number of non-hydrogen atoms | 13931 | 66334 | 13261 |
| macromolecules | 13191 | 6590 | 13180 |
| ligands | 0 | 0 | 10 |
| solvent | 740 | 44 | 71 |
| Protein residues | 1729 | 866 | 1728 |
| RMS(bonds) | 0.011 | 0.017 | 0.015 |
| RMS(angles) | 1.63 | 2.25 | 2.14 |
| Ramachandran favored (%) | 97.14 | 88.46 | 89.37 |
| Ramachandran allowed (%) | 2.69 | 9.32 | 8.29 |
| Ramachandran outliers (%) | 0.18 | 2.21 | 2.34 |
| Rotamer outliers (%) | 3.15 | 18.87 | 12.84 |
| Clashscore | 3.42 | 14.9 | 10.48 |
| Average B-factor | 35.98 | 84.38 | 69.03 |
| macromolecules | 35.77 | 84.8 | 69.17 |
| solvent | 39.81 | 22.41 | 45.74 |
| TLS groups | None | 4 | 8 |

**Supplementary table 7.** Crystallographic data collection and refinement statistics. Statistics for the highest resolution shell are shown in parentheses. The first two column datasets were treated with anisotropic data inclusion criteria, for which parameters of the ellipsoid are described.
